# Accuracy of multiple sequence alignment methods in the reconstruction of transposable element families

**DOI:** 10.1101/2021.08.17.456740

**Authors:** Robert Hubley, Travis J. Wheeler, Arian F.A. Smit

## Abstract

The construction of a high-quality multiple sequence alignment (MSA) from copies of a transposable element (TE) is a critical step in the characterization of a new TE family. Most studies of MSA accuracy have been conducted on protein or RNA sequence families where structural features and strong signals of selection may assist with alignment. Less attention has been given to the quality of sequence alignments involving neutrally evolving DNA sequences such as those resulting from TE replication. Such alignments play an important role in understanding and representing TE family history. Transposable element sequences are challenging to align due to their wide divergence ranges, fragmentation, and predominantly-neutral mutation patterns. To gain insight into the effects of these properties on MSA accuracy, we developed a simulator of TE sequence evolution, and used it to generate a benchmark with which we evaluated the MSA predictions produced by several popular aligners, along with Refiner, a method we developed in the context of our RepeatModeler software. We find that MAFFT and Refiner generally outperform other aligners for low to medium divergence simulated sequences, while Refiner is uniquely effective when tasked with aligning high-divergent and fragmented instances of a family. As a result, consensus sequences derived from Refiner-based MSAs are more similar to the true consensus.

## Introduction

The ongoing explosion in the number of sequenced organisms highlights the need for reliable and thorough automated genome annotation pipelines. Most of the vertebrate genome finds its ultimate origin in transposable elements (TEs)(1–5), which have an enormous impact on genome activity and evolution(6–9). Due to the volume and importance of TEs, complete annotation of genomes depends on accurate identification and modeling of TE families (10). A central aspect of that process is the gathering of instances of each family, and the creation of multiple sequence alignments (MSAs) of those instances; these MSAs are used to derive a consensus sequence (1, 11–13) and optionally a profile hidden Markov model (HMM) (14) for each family. An accurate family-level TE MSA is critical for sensitive annotation of genomic copies, precise classification of TE families, reconstruction of encoded proteins, and family age estimation; this motivates an intense interest in the accuracy of methods producing MSAs for these sequence families.

Computational approaches to MSA seek to optimize one of several scoring models, and optimal solution of commonplace models is computationally intractable (15). Over the years, a multitude of MSA tools has been developed, each employing its own set of heuristics for achieving good alignment speed. The combination of heuristic and scoring functions leads to varying alignment accuracy. MSA of TE instances poses unique challenges, in that these instances can exhibit high sequence divergence, are often highly fragmented, and are dominated by neutral mutation patterns. Here, we seek to evaluate the efficacy of several commonly used tools in recovering accurate MSAs of neutrally evolving fragments of transposable element sequences.

TEs are prodigious generators of repetitive sequences in most genomes; their relationships can be difficult to recover due to rapid lineage bursts, complex recombination histories, and high rates of neutral mutation. The generation of a MSA from copies of a TE family is an important step in reconstructing the ancestral state of the TE and generating sequence models for genome annotation(16). TE sequences complicate alignment in several important ways: (1) Instances are often fragmented due to poor insertion fidelity, large deletions, or interruptions by insertions of other TEs. (2) Due to their mostly neutral decay, there are generally no conserved regions that can anchor the alignment or open reading frames free from indel accumulation. (3) Copies are often derived from a TE that was rapidly evolving in a genome; therefore they represent a mixture of ancestral forms. (4) Low complexity regions and internal repetition are common features. (5) The oldest detectable TE copies have accumulated over 35% substitutions since their arrival and given their neutral decay, have a substitution level of more than 70% between each other at the DNA level.

Evaluation of MSA tool accuracy usually depends on protein benchmarks, consisting of real sequences (17–20), based on structural (PREFAB(21)) or hand-curated alignments(e.g. BAliBASE(22), SABmark(23), HomFam(24), HOMSTRAD(25), and OXBENCH(26)). BRaliBase(27)) stands out as a rare benchmark for nucleotide (RNA) alignment. Simulated sequence evolution datasets have also been used to evaluate MSA tools(28–31), providing the means to produce larger test sets, a wide range of sequence divergence characteristics, and supporting the generation of DNA-specific benchmarks. Missing from these analyses are assessments of accuracy for MSAs of neutrally diverged sequences, such as ancient copies of TEs.

Sequence simulation tools have themselves evolved over time. Early efforts focused primarily on the generation of sequences along a fixed phylogeny, allowing for mutations based on time reversible substitution models(32–35). These led to more sophisticated evolvers with support for insertion and deletion (indels) mutations, empirical substitution matrices, and branch dependent mutation rates(36–42). Context dependent mutation rates have also been developed in some simulators; for instance the Evolver package(43) models special mutation rates for highly mutagenic CpG dinucleotides, and Trevolver(44) implements a triplet substitution model that accounts for first-order flanking effects.

To the best of our knowledge, no formal evaluation of MSA tools has been previously conducted in the context of TE sequence families. To study this, we developed a new sequence simulator (TEForwardEvolve), and used it to generate a benchmark of simulated MSAs. Using this benchmark, supplemented with a collection of hand-curated mammalian TE MSAs, we evaluated several current MSA tools: MUSCLE(21), MAFFT(45), Dialign-TX(46), Kalign(47), FSA(48), Clustal Omega(49). We additionally considered Refiner(50), a transitive alignment method that we developed for our RepeatModeler software package, and which we have employed in the reconstruction of TE families for many years. We demonstrate that MAFFT generally outperforms other generic alignment tools, and that our Refiner method produces comparable results for low-divergence sequences, and superior alignments when confronted with high levels of sequence fragmentation and sequence divergence.

## Results

### Simulated Trees and Sequences

Phylogenetic trees approximating the evolutionary patterns of DNA transposons and LINE families were randomly generated using a custom-made tool. The relationship of DNA transposon copies (Figure 1A) is typically a (near) star phylogeny, with any branching occurring randomly and early on. This reflects the fact that, due to random selection of the genomic template by the transposase, class II transposons tend to exhibit a short burst of activity before going extinct (1, 51) and leave many neutrally decaying copies in the genome. DNA transposon star phylogenies may be contrasted with those of LINEs (Figure 1B), in which most copies are derived from a single dominant lineage of LINE TEs (the so-called master-gene model of evolution (52, 53)); the resulting phylogenies approach those of pseudogenes of a rapidly evolving cellular gene.

**Figure 1:**
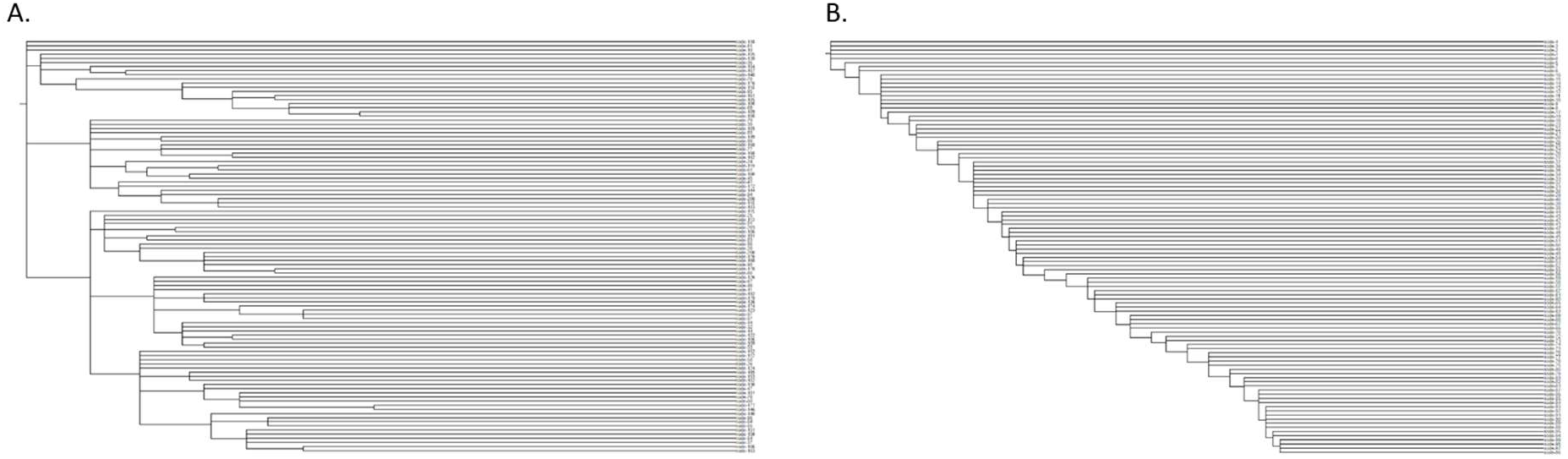
Phylogenetic Trees For Simulations. (A) Random template phylogeny generated typical of DNA transposons, and (B) master gene phylogeny for LINE families.

Sequences were simulated along these trees using a forward evolution sequence simulator seeded with a class-specific TE consensus sequence (See Methods for details). The DNA transposon sequence simulation was seeded with the Tigger1 family consensus(54), and the LINE tree was seeded with the L2 consensus(1). Simulation was run with ten replicates at 18 evolutionary time increments, producing 180 simulated sequence sets and reference MSAs. Evaluation with Charlie1 (DNA transposon) and CR1 (LINE) produced similar results, and are presented in the supplementary material.

### Alignment Reconstruction Accuracy

For each alignment method evaluated, a predicted MSA was produced from each sequence set and compared to the simulated MSA using the Sum-of-Pairs Score (SPS, aka developer score)(55, 56). SPS measures the fraction of residue pairs sharing a column in the reference MSA that also share a column in the predicted MSA; it provides a global assessment for the accuracy of the prediction.

We computed SPS for all methods over a wide range of sequence divergence and for both TE classes (Figure 2). The Kruskal-Wallis H test was performed over the replicates to assess the significance of the distribution of score means and confirmed a significant separation of SPS for both the DNA transposon simulations (p-value=4.17e-24) and the LINE simulation (p-value=2.15e-31). To compare relative performance of each pair of methods, we used a Wilcoxon signed rank test and determined that MAFFT and Refiner significantly outperformed other methods (see Supplementary material for the full pairwise comparison table).

**Figure 2:**
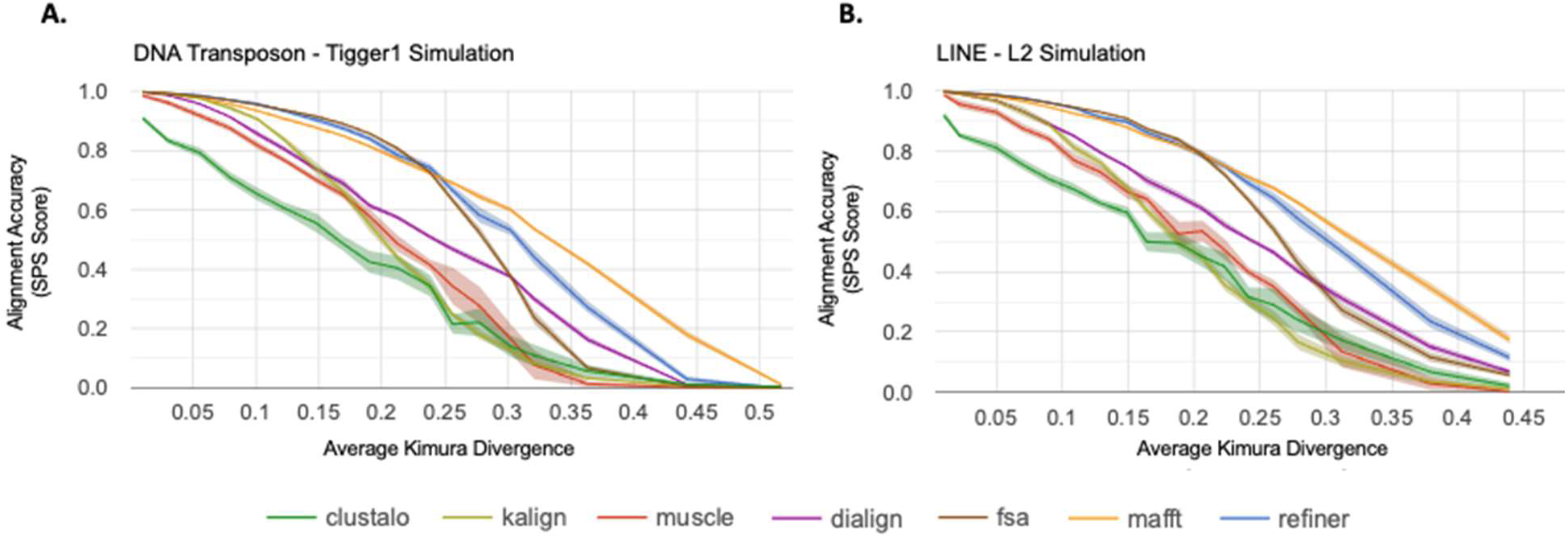
MSA accuracy with respect to sequence divergence. The MSA alignment accuracy for each method assessed using the sum-of-pairs (SPS) score over a wide range of sequence divergence. For each tool/divergence combination, 10 replicates were performed. The center line of each band shows the mean SPS score for the tool, while the surrounding shaded region shows the 95% confidence interval. (A) The Tigger1 DNA transposon family average SPS scores over 10 replicates. (B) The L2 LINE family average SPS scores over 10 replicates.

### Alignment as a Basis for Consensus Sequences

Consensus sequences can be derived from MSAs by choosing the most likely ancestral base at each position in the MSA (considering only positions with high occupancy)(57); this has, for example, long been the source of family consensus sequences used in annotating TEs. When TE copies form a star phylogeny, which appears to be the case for most class II transposon copies in mammals(58), the consensus will be identical to the ancestral sequence of the active TE. In case of the LINE MSA, a consensus may approach an average of the evolving active TE.

To compare the accuracy of consensus reconstruction we define a new score metric. Let C_p_ be the consensus sequence derived from the predicted MSA, and C_r_ the consensus sequence produced from the reference MSA. To measure similarity of C_p_ to C_r_, we aligned them to each other via Needleman-Wunch global pairwise alignment(59) C_p_ ∼ C_r_ (see Methods), and aligned C_r_ to itself to produce alignment C_r_ ∼ C_r_. If the predicted MSA is perfectly accurate, C_p_ will be identical to C_r_, so that score(C_p_ ∼ C_r_) == score(C_r_ ∼ C_r_). An inaccurate predicted MSA will cause score(C_p_ ∼ C_r_) < score(C_r_ ∼ C_r_). Figure 3 shows the extent to which computed alignments support recovery of accurate consensus sequences, by presenting the fraction of the ideal score that is lost with the predicted alignment: (score(C_r_ ∼ C_r_) - score(C_p_ ∼ C_r_)) / score(C_r_ ∼ C_r_). In the case of an extremely poor predicted MSA, the predicted consensus may lead to a negative score(C_p_ ∼ C_r_), so that the lost score exceeds the ideal score (i.e. more than 100% of the score is lost). This score-loss measure is captured for each tool at a variety of divergence levels.

**Figure 3:**
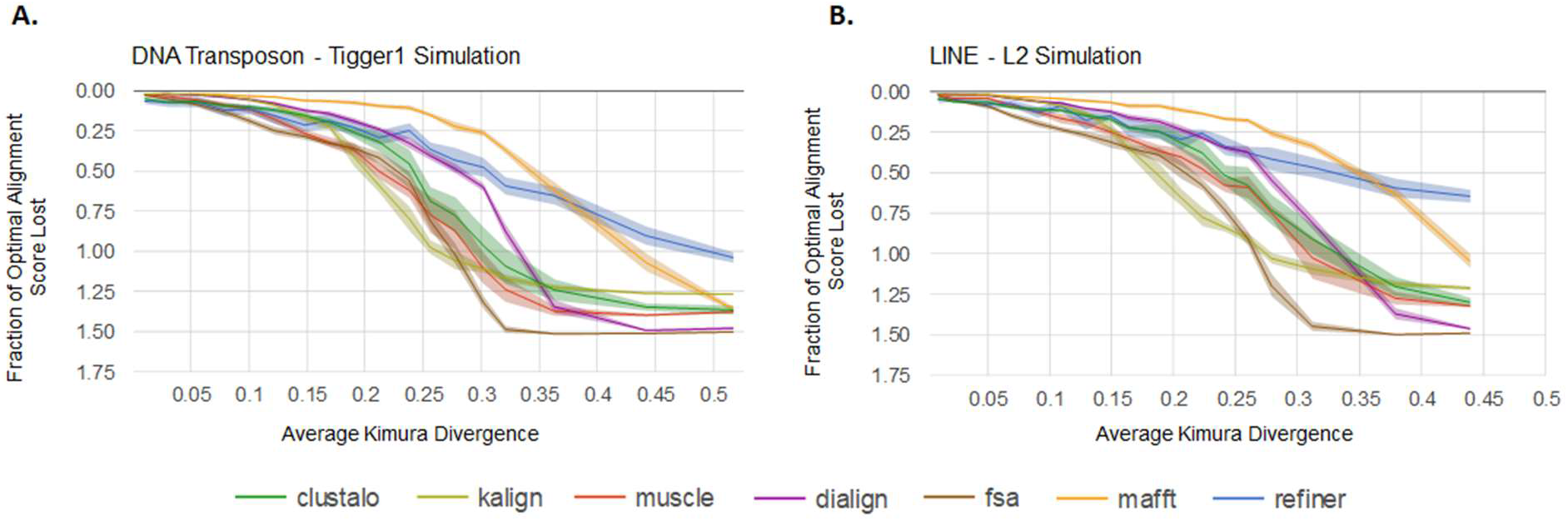
Accuracy of derived consensus sequence with respect to sequence divergence. Comparison of the predicted MSA-based consensus with the reference MSA-based consensus,for a range of sequence divergences.. (A) For simulated Tigger1 sequences with variable levels of Kimura divergence, this plot shows the fraction (score(C_r_ ∼ C_r_)-score(C_p_ ∼ C_r_))/ score(C_r_ ∼ C_r_), which corresponds to how effective the computed MSA is at producing a consensus sequence (C_p_) that agrees with the one for the simulated sequence (C_r_). Scores are for the Needleman-Wunsch (NW) global alignment algorithm (see methods). (B) The same fraction-of-optimal-score is captured, but for sequences simulated from L2. Center line of each band shows the mean loss of score for each tool, while the surrounding shaded region shows the 95% confidence interval.

### Effect of Sequence Fragmentation

We evaluated the effect of fragmentation on MSA reconstruction as above, comparing the predicted MSA to the reference MSA, assessing both low divergence (1% avg Kimura divergence(60)) and high divergence (28% avg Kimura) sequences. The mean fragmentation size was varied from 75-1200bp based on observed fragmentation patterns in mammalian TE copies (see supplemental materials for details). Figure 4 shows the effects of fragmentation level on SPS and provides a visualization of the patterns for the fragmentation extremes. Most tools performed well on low-divergence sequences over a wide range of fragment sizes; at higher sequence divergence and fragmentation, Refiner outperforms all methods tested (Wilcoxon p-value<=1.1e-11), with MAFFT, Dialign, and FSA outperforming the rest. We also explored the effect of fragmentation on the accuracy of MSA-derived consensus sequences, as in the previous section (Figure 5). At high sequence divergence levels, Refiner was the only tool with partial retention of correct alignment score, showing that it effectively produces MSAs that yield accurate consensus sequences.

**Figure 4:**
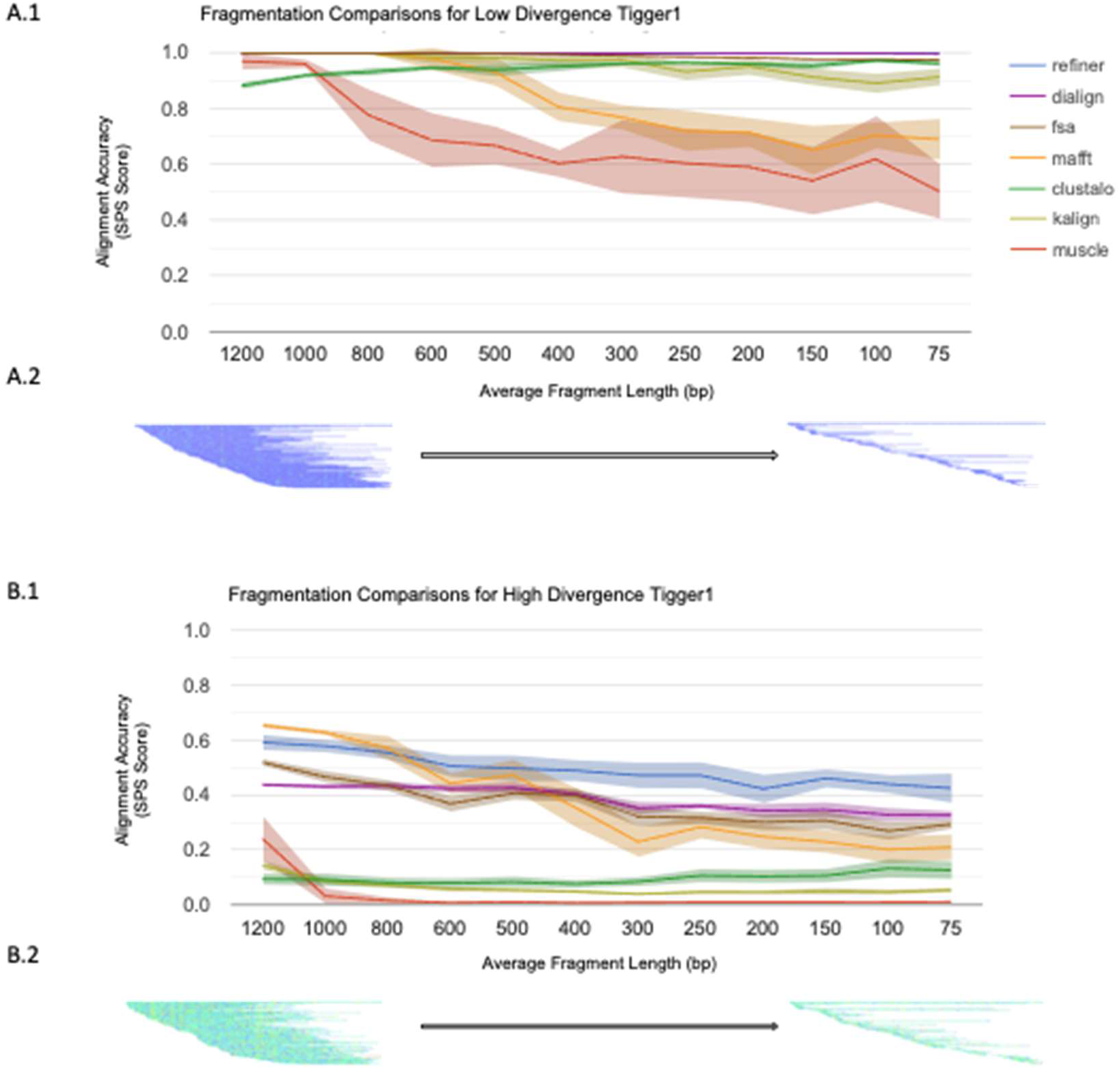
MSA accuracy with respect to sequence fragmentation. (A.1) The SPS results for simulations of Tigger1 (1% Kimura div) over increasing levels of sequence fragmentation. Fragment lengths are sampled from a distribution around the given mean (x-axis) with a standard deviation of 300. (A.2) A visualization of the fragmentation of the reference MSA for the least fragmented and most fragmented datasets. Each line represents a single fragment; warmer colors represent higher sequence divergence over 10bp windows in the alignment. (B.1) The SPS results for simulations of Tigger1 (28% Kimura div) over increasing levels of sequence fragmentation, with fragment length sampled as above. (B.2) Heatmap visualization of the fragmented MSA, as with (A.2), but for the higher divergence Tigger1 benchmark. Center line of each band shows the mean SPS for each tool, while the surrounding shaded region shows the 95% confidence interval.

**Figure 5:**
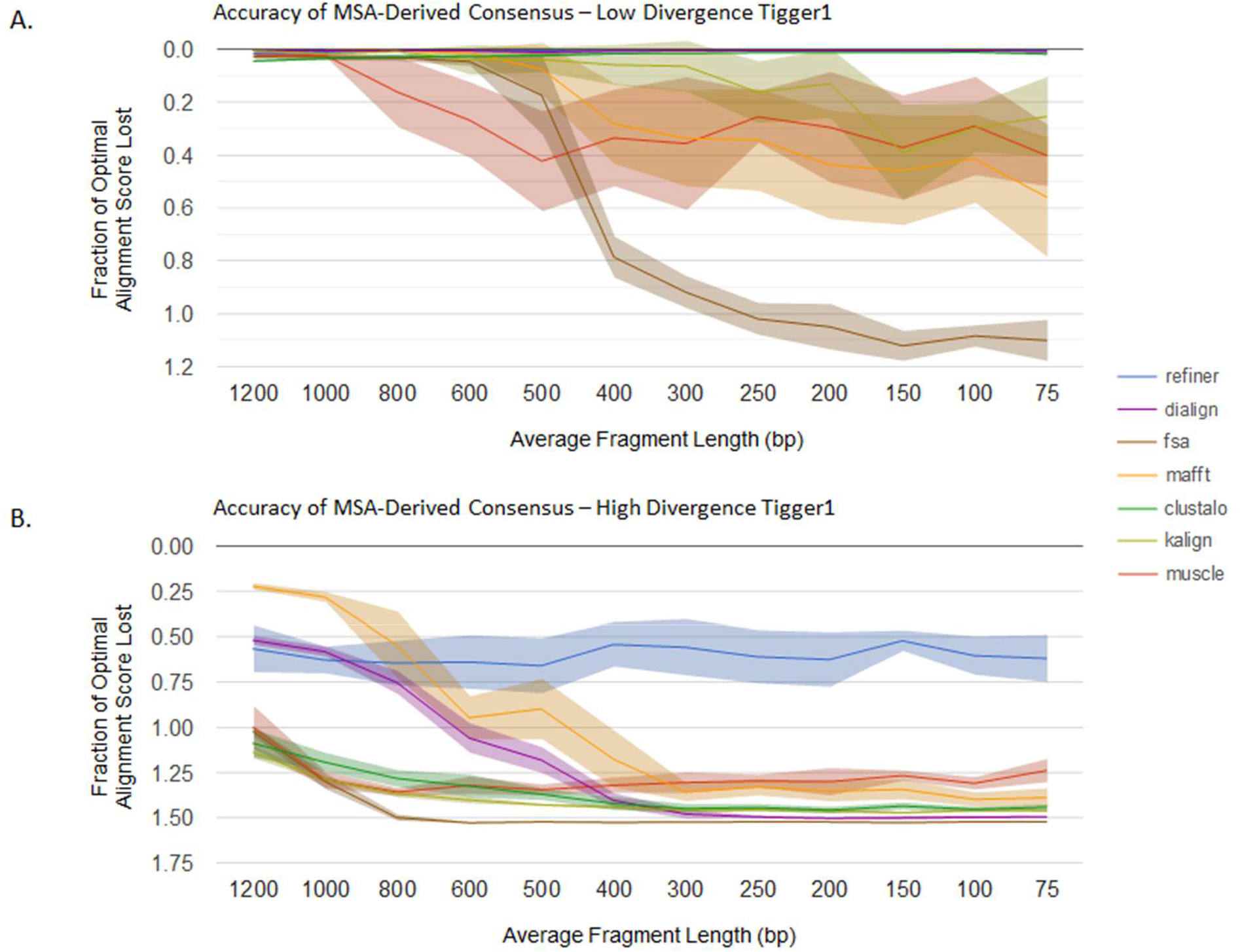
Accuracy of derived consensus with respect to sequence fragmentation. Comparison of the predicted MSA-based consensus with the reference MSA-based consensus, for fragmented sequences. (A) Fraction of ideal sequence alignment score, as in Figure 3; input sequences are low-divergence fragments (Avg Kimura div = 1%) as from Figure 4A. (B) Same as in A, but with high-divergence fragment inputs (Avg Kimura div = 28%) as from Figure 4B.

### Comparison with Natural Sequence

We selected four DNA transposons: Zaphod(61), Zaphod2(62), Tigger10(63), and Arthur2(64) to compare reconstruction accuracy at the protein level. These were selected for their age (Tigger10 and Arthur2 predate our speciation from marsupials) and the high divergence of human copies to each other are expected to form a challenge for MSA tools. Our manually created consensus sequences, aided by copies from reconstructed ancestral mammalian genomes(65) have full open reading frames for transposases. As DNA transposons, they are expected to have a star phylogeny, so that an accurate MSA should recreate the ORFs of the active elements. For each family, 150 annotated instances were aligned with each tool. A consensus was generated from each predicted MSA, then blastx (default: matrix=blosom62, E=10.0, gap_open=11, gap_ext=1) was used to compare the consensus to the protein. The alignment bit scores for the top hits are shown in Table 1. Only consensus sequences derived from the refiner MSAs had matches for each element. These matches were also much stronger.

**Table 1:**
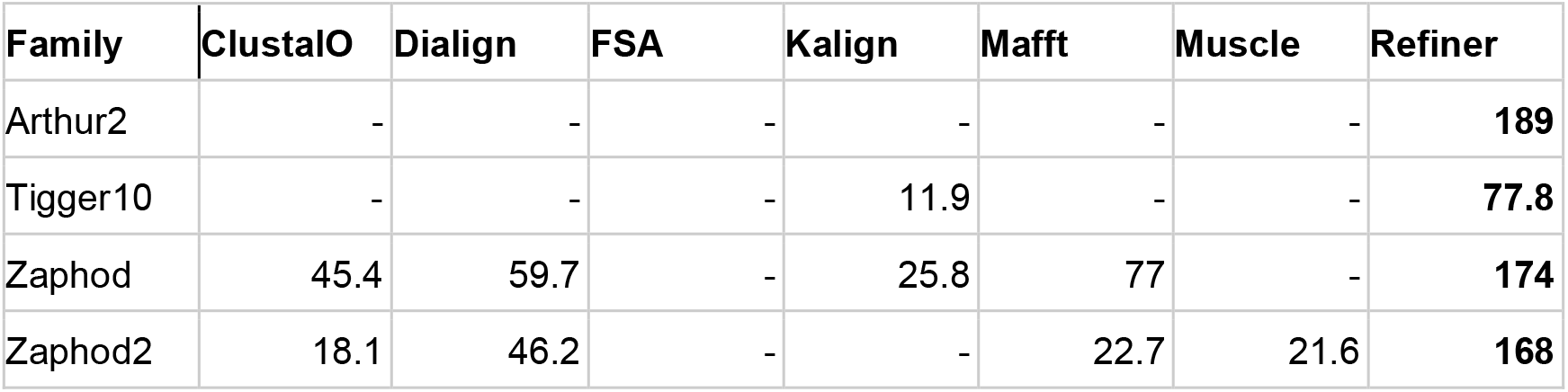
Protein reconstruction assessment for four mammalian TE families using 150 human-derived copies. A consensus model was built from each MSA and compared to the curated protein sequence using blastX (matrix: BLOSUM62). The highest blastx bit score is recorded for each aligner; a value of “-” indicates that no alignment could be found.

## Discussion

Using our new method TEForwardEvolve, we simulated the evolution of TE families over a wide variety of sequence divergence and fragmentation to investigate the impact on MSA prediction. The evaluated tools exhibited similar patterns of performance degradation for both TE classes as the sequence divergence was increased; with statistically significant performance differences among them. For full-length sequence inputs,MAFFT and Refiner maintained the highest alignment accuracy over the range of divergences.

SPS (aka the developer score) is an established and generally informative measure of global MSA reconstruction; however, it doesn’t appear to fully reflect the performance of MSA tools in the context of consensus sequence prediction. This can be seen most clearly in the performance of Refiner at high sequence divergences, where it is outcompeted by MAFFT’s SPS score while simultaneously outcompeting MAFFT when evaluating the accuracy of the MSA-derived consensus sequence. We hypothesize that local alignment could be key to explaining the difference. The consensus is negatively influenced by the tendency of many MSA tools to force mismatched sequence regions together in the MSA (66, 67); local alignment avoids this at the expense of a complete alignment. In addition, short regions of misalignment in an MSA, while not penalized heavily by the SPS metric, can lead to incorrect consensus generation. With highly fragmented sequences, FSA occasionally produced alignments where a majority of the sequences were incorrectly anchored at the start of the alignment for a short stretch (3bp) followed by large gaps before continuing in roughly the correct location in the overall MSA (See supplemental). In cases such as this, the alignment artifact would cause a consensus caller to consider the sequences contained in the gap as probable insertions and generate a dramatically shortened consensus.

Our results suggest that even low levels of fragmentation, when combined with higher sequence divergence, provides a significant challenge to MSA tools. Faced with fragmentary input, Refiner produced MSAs that were significantly more accurate than other tools, highlighting the value of its iterative transitive alignment approach for this challenging form of input. For this study, fragmentation is simulated as a random process, whereas fragmentation processes exhibit biases for many TE classes (e.g. 5’ truncation in LINE elements, or the generation of solo LTRs through recombination). We plan to extend our simulator, TEFowardEvolve, to explore the effect of these fragmentation architectures on MSA prediction as well as additionally consider SINE and LTR TE classes.

Finally, simulation results were validated with an evaluation using manually curated protein sequences and instances of the TE family sequences that encode them. Only one tool (Refiner) was consistently successful in computing an MSA-based consensus that matched the corresponding protein in a blastx search. The classification of a TE family is greatly facilitated by comparing to existing TE protein databases, underscoring the importance of an accurate MSA reconstruction.

## Materials and Methods

### Tree Generation

A custom-made tool (genRandomTETrees.pl) was used to simulate phylogenetic trees for DNA Transposon and LINE TE families. For DNA Transposons, the tree is expanded by randomly choosing a parent node from the existing tree and appending a new child node with a random branch length between 0-10. Once the target number of extant nodes has been reached (100), the post-extinction phase of the sequence lifecycle is simulated by adjusting extant (leaf) nodes’ branch lengths to reach the target root-to-leaf length (100). Branch lengths in the tree do not equate to a specific unit of time; rather, they establish the relative duration of each branch. For the purposes of sequence simulation, the notional duration of a branch (i.e. the amount of mutation) is controlled by the simulator parameter “generations per unit time” (GPUT).

To simulate LINE phylogenies, the tree is expanded by adding a randomly determined number of children (2-5) with randomly chosen branch lengths (0-5) to the current parent node (initially the root of the tree). One of the new children is randomly picked to be the new parent node (or “master gene”) and the process is iterated until the target number of extant nodes is reached (100). The branch lengths for extant nodes are then adjusted in a similar manner to the DNA transposon trees.

### Sequence Simulation

While there are many sequence evolution simulators currently available, none provide all the features necessary for realistic TE sequence simulation in one package: nucleotide simulation, indel simulation, context dependent (trinucleotide) substitution matrices, and fragmentation simulation. Inspired by the release of TRevolver(44), which supports tri-nucleotide substitution context, we developed TEForwardEvolve, which supplements tri-nucleotide substitution with simulation of indels and fragmentation.

For each class of TE, the simulator is provided a prototypical TE sequence (Tigger1/Charlie1 for DNA transposons and L2/CR1 for LINEs), a simulated phylogenetic tree, a context-dependent substitution rate matrix, indel parameters, the number of generations represented in the tree, and fragmentation parameters. The substitution rate matrix consists of all triplet pairs where the center base is allowed to change and the edge bases provide 1bp of flanking context (64×64 matrix); rates were derived from a study of 160,000 non-coding sites in a set of mammalian genomes(68). Indel lengths were modeled using a power law (Zipfian) probability distribution (insertion/deletion mean length = 1.7, insertion/deletion max length = 20) with an occurrence rate of 0.20 (insertion rate = 0.08, deletion rate = 0.12) relative to an average substitution rate of 1. Finally, fragmentation is simulated by selecting fragment sizes from a log normal distribution, a minimum fragment size and a randomly chosen start position to select only a portion of the parent sequence for duplication. Optionally, a minimum number of full-length sequences may be set such that fragmentation begins only after the minimum number of full-length sequences has been generated.

To study the impact of sequence substitution level, TEFowardEvolve was run with increasing values for the generations-per-unit-time (GPUT) simulation parameter (100-6000). This translates to an average Kimura divergence range of 1-52% for Tigger1 and 1-44% for L2 simulations. To study the impact of fragmentation, TEForwardEvolve was run with increasing levels of fragmentation (mean copy lengths ranging from 1200 down to 75, with a standard deviation of 300, minimum fragment size=50, minimum full-length sequences=2) at two substitution levels (GPUT 100, 3000). For each parameterization, 10 replicate simulations were run.

### Refiner Methodology

Our Refiner method works by establishing a single template sequence, aligning all sequences to that template, then producing a MSA based on the way that those sequences align to the template (in what we call a transitive alignment). In the first pass, the template is chosen from the input sequences by picking the sequence with the best cumulative pairwise alignment score to all other sequences. The resulting MSA is used to produce a consensus, which is used as the template for iterative rounds of transitive alignment, in the form of Expectation Maximization. This process is repeated until convergence; see (50, 69) for details. The final reference sequence is the consensus for the family.

In this local-alignment strategy, characters that do not align to the template sequence are either arbitrarily aligned to each other (internal insertions) or not included in the alignment at all. This is not a problem in the context of RepeatModeler, since these are not part of high occupancy columns, so not part of the final consensus. For the purpose of MSA SPS evaluation, all characters must be present in the final alignment, so we simply add all such characters to the MSA such that they are not aligned to any other character.

The consensus caller used by Refiner employs two stages. The caller initially identifies the highest scoring character (‘A’,’C’,’G’,’T’,’N’ or ‘-’) for each column from the subset of sequences aligning over it. The first step uses a matrix that reflects observed neutral DNA substitution patterns and is similar to matrices developed for RepeatMasker. For organisms with CpG methylation, resulting in high turnover of CG to CA and TG, a second pass over all dinucleotides in the initial consensus sequence are evaluated for reassignment to ‘CG’ by registering the frequency of the most common products of CpG mutation, aligned CA and TG dinucleotides(69).

### MSA Evaluation

For each sequence evolution simulation, TEForwardEvolve provides a reference MSA for comparison to the ones predicted by the alignment tools. The qscore (21) tool was used to compute various metrics on the predicted MSAs including SPS, which is the fraction of aligned residue pairs in the reference alignment that are correctly aligned in the predicted alignment.

The quality of the predicted MSA was also assessed by the quality of the consensus it produces (see previous section for a description of the consensus caller). As a measure of consensus quality, we computed the Needleman-Wunsch global alignment score for an alignment of the predicted MSA (C_p_) to the reference MSA (C_r_), and for an alignment of C_r_ to itself, then computed the fraction of the ideal score (for C_r_ ∼ C_r_) that is lost in the alignment C_p_ ∼ C_r_: (score(C_r_ ∼ C_r_) - score(C_p_ ∼ C_r_)) / score(C_r_ ∼ C_r_). Alignment was performed with a custom scoring matrix and gap parameterization described in the supplemental materials.

### MSA Tools and Parameters

The MSA tools covered in this evaluation are shown in Table 2, including versions and any non-default parameter settings. No attempt was made to optimize parameters aside from ensuring that DNA specific defaults were used. For Kalign, the DNA/RNA default parameters were based on instructions on the software website (https://msa.sbc.su.se/cgi-bin/msa.cgi). For MAFFT, the linsi algorithm was specifically chosen for evaluation based on expectation that this will provide greatest accuracy.

**Table 2:**
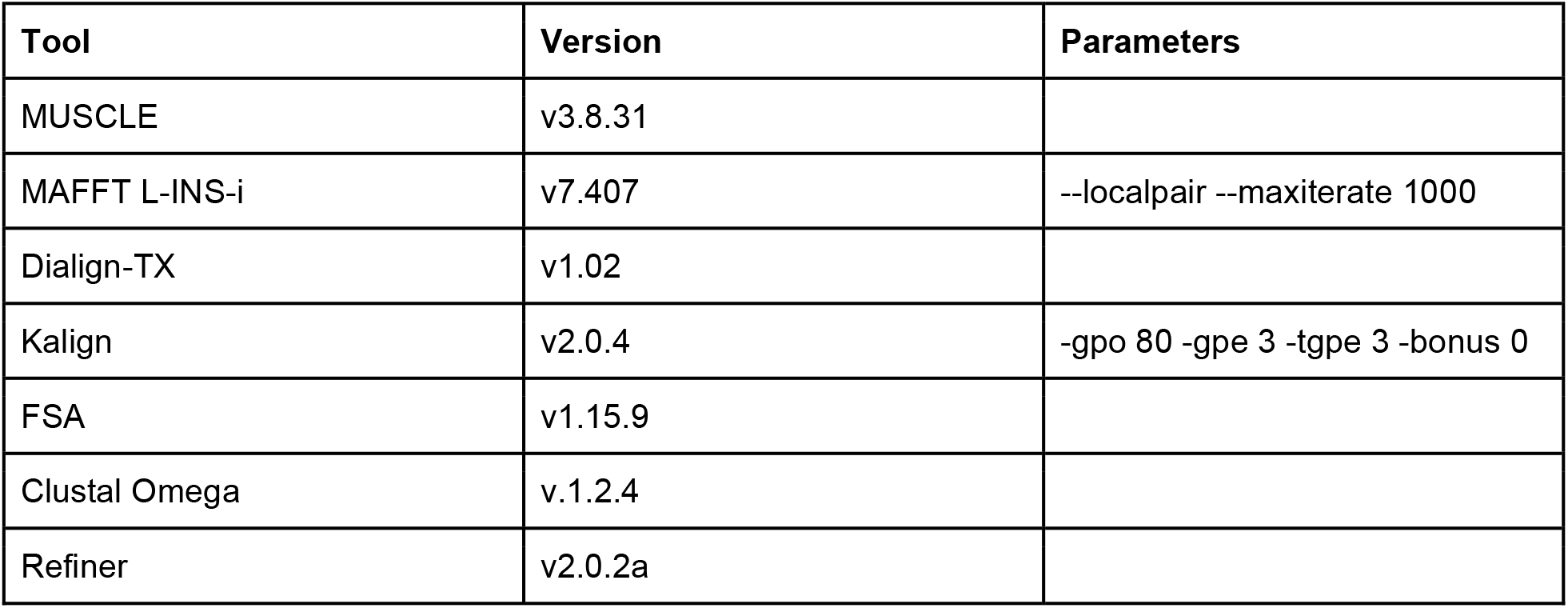
MSA Software evaluated. The version of each tool evaluated and any specific parameters provided.

## Availability

TEForwardEvolve, genRandomTETrees, analysis scripts, Nextflow (70) job management scripts, along with simulated data generated as part of this study, are available on github at: https://github.com/Dfam-consortium/TEForwardEvolve, release tag “1.0-paper”. MSA tool results, graphs and alignment visualizations may be downloaded from: https://www.dfam.org/web_download/Publications/MSABench2021.

## Acknowledgements

We would like to thank Adam Siepel for sharing his estimated triplet rate matrices with us. We would also like to acknowledge Paul Edlefsen for the many helpful discussions on the topic.

## Author Contributions

RH wrote the sequence simulation software, conducted the MSA analysis, and wrote the draft of the manuscript. TJW and AFS provided feedback on methods and edited the manuscript.

