## Supplemental Data for "Accuracy of multiple sequence alignment methods in the reconstruction of transposable element families"

#### Kruskal-Wallis One Way Analysis of Variance and Wilcoxon Signed Rank Post-hoc Tests

##### Sum of Pairs (SPS) Metrics

For each MSA tool evaluated, the full set of 180 SPS measurements (10 replicates, 18 divergence bins) were collected. A Kruskal-Wallis H-test was performed to assess if there is a significant difference in the accuracy of alignments between each pair of MSA tools. Additionally, a Wilcoxon signed rank post-hoc test was performed to generate multiple comparison tables for all pairs of methods. Section S1 contains the tables for the data presented in the main paper while section S2 includes tables for additional tree simulations, and additional seed sequences.

##### Section S1

SPS Statistics for paper-data/DNATransTree-1-Tigger1-R3S-eval/replicates.csv
- Calculated on all replicate-parameter values
- mean_diff is the difference in mean SPS scores; for example, the mean SPS of refiner is 0.19 higher than the mean SPS of muscle

Kruskal-Wallis H-test
H=122.8175068514761, p=4.1695554830026675e-24

Wilcoxon signed rank test: mean_diff [p-val]

| | refiner | muscle | mafft | dialign | kalign | fsa | clustalo |
|---|---|---|---|---|---|---|---|
| refiner | | 0.19 [7.0e-31] | -0.02 [7.6e-02] | 0.12 [7.8e-30] | 0.19 [7.8e-29] | 0.03 [8.6e-06] | 0.28 [1.2e-30] |
| muscle | -0.19 [7.0e-31] | | -0.21 [2.7e-31] | -0.07 [1.9e-29] | -0.00 [1.5e-01] | -0.16 [6.8e-31] | 0.09 [4.1e-20] |
| mafft | 0.02 [7.6e-02] | 0.21 [2.7e-31] | | 0.14 [1.2e-30] | 0.21 [5.0e-29] | 0.06 [1.8e-02] | 0.30 [2.8e-31] |
| dialign | -0.12 [7.8e-30] | 0.07 [1.9e-29] | -0.14 [1.2e-30] | | 0.07 [8.1e-10] | -0.08 [4.3e-17] | 0.16 [3.3e-30] |
| kalign | -0.19 [7.8e-29] | 0.00 [1.5e-01] | -0.21 [5.0e-29] | -0.07 [8.1e-10] | | -0.16 [9.3e-28] | 0.09 [2.4e-15] |
| fsa | -0.03 [8.6e-06] | 0.16 [6.8e-31] | -0.06 [1.8e-02] | 0.08 [4.3e-17] | 0.16 [9.3e-28] | | 0.24 [6.8e-29] |
| clustalo | -0.28 [1.2e-30] | -0.09 [4.1e-20] | -0.30 [2.8e-31] | -0.16 [3.3e-30] | -0.09 [2.4e-15] | -0.24 [6.8e-29] | |

SPS Statistics for paper-data/DNATransTree-1-Tigger1-R3S-gput100-mfl2-eval/replicates.csv
- Calculated on all replicate-parameter values

Kruskal-Wallis H-test
H=454.07361102460294, p=6.518631485795493e-95

Wilcoxon signed rank test: mean_diff [p-val]

| | refiner | muscle | mafft | dialign | kalign | fsa | clustalo |
|---|---|---|---|---|---|---|---|
| refiner | | 0.32 [2.0e-21] | 0.17 [8.9e-17] | 0.00 [1.9e-14] | 0.04 [4.6e-19] | 0.01 [2.5e-16] | 0.05 [2.0e-21] |
| muscle | -0.32 [2.0e-21] | | -0.15 [2.8e-13] | -0.32 [2.9e-21] | -0.28 [1.6e-20] | -0.31 [2.0e-21] | -0.27 [8.0e-17] |
| mafft | -0.17 [8.9e-17] | 0.15 [2.8e-13] | | -0.17 [6.2e-14] | -0.13 [3.2e-13] | -0.16 [1.6e-15] | -0.12 [1.0e-09] |
| dialign | -0.00 [1.9e-14] | 0.32 [2.9e-21] | 0.17 [6.2e-14] | | 0.04 [8.7e-15] | 0.01 [8.3e-12] | 0.05 [2.0e-21] |
| kalign | -0.04 [4.6e-19] | 0.28 [1.6e-20] | 0.13 [3.2e-13] | -0.04 [8.7e-15] | | -0.03 [6.8e-11] | 0.01 [2.3e-02] |
| fsa | -0.01 [2.5e-16] | 0.31 [2.0e-21] | 0.16 [1.6e-15] | -0.01 [8.3e-12] | 0.03 [6.8e-11] | | 0.04 [1.4e-20] |
| clustalo | -0.05 [2.0e-21] | 0.27 [8.0e-17] | 0.12 [1.0e-09] | -0.05 [2.0e-21] | -0.01 [2.3e-02] | -0.04 [1.4e-20] | |

SPS Statistics for paper-data/DNATransTree-1-Tigger1-R3S-gput3000-mfl2-eval/replicates.csv
- Calculated on all replicate-parameter values

Kruskal-Wallis H-test
H=669.0718192551245, p=2.90637951578182e-141

Wilcoxon signed rank test: mean_diff [p-val]

| | refiner | muscle | mafft | dialign | kalign | fsa | clustalo |
|---|---|---|---|---|---|---|---|
| refiner | | 0.46 [2.0e-21] | 0.11 [1.1e-11] | 0.11 [2.0e-20] | 0.43 [2.0e-21] | 0.13 [3.8e-21] | 0.39 [2.0e-21] |
| muscle | -0.46 [2.0e-21] | | -0.35 [2.0e-21] | -0.35 [2.0e-21] | -0.03 [1.7e-12] | -0.33 [2.0e-21] | -0.07 [1.0e-12] |
| mafft | -0.11 [1.1e-11] | 0.35 [2.0e-21] | | -0.01 [5.3e-01] | 0.31 [2.0e-21] | 0.01 [3.5e-01] | 0.28 [5.4e-21] |
| dialign | -0.11 [2.0e-20] | 0.35 [2.0e-21] | 0.01 [5.3e-01] | | 0.32 [2.0e-21] | 0.02 [4.5e-05] | 0.29 [2.0e-21] |
| kalign | -0.43 [2.0e-21] | 0.03 [1.7e-12] | -0.31 [2.0e-21] | -0.32 [2.0e-21] | | -0.30 [2.0e-21] | -0.04 [1.1e-11] |
| fsa | -0.13 [3.8e-21] | 0.33 [2.0e-21] | -0.01 [3.5e-01] | -0.02 [4.5e-05] | 0.30 [2.0e-21] | | 0.27 [2.0e-21] |
| clustalo | -0.39 [2.0e-21] | 0.07 [1.0e-12] | -0.28 [5.4e-21] | -0.29 [2.0e-21] | 0.04 [1.1e-11] | -0.27 [2.0e-21] | |

SPS Statistics for paper-data/LINETree-1-L2-R3S-gput100-mfl2-eval/replicates.csv
- Calculated on all replicate-parameter values

Kruskal-Wallis H-test
H=551.6357511140836, p=6.26877337556657e-116

Wilcoxon signed rank test: mean_diff [p-val]

| | refiner | muscle | mafft | dialign | kalign | fsa | clustalo |
|---|---|---|---|---|---|---|---|
| refiner | | 0.50 [2.0e-21] | 0.21 [3.9e-18] | 0.00 [2.2e-13] | 0.13 [2.3e-20] | 0.01 [5.7e-17] | 0.04 [2.0e-21] |
| muscle | -0.50 [2.0e-21] | | -0.28 [3.8e-21] | -0.50 [2.0e-21] | -0.37 [2.4e-21] | -0.49 [2.0e-21] | -0.46 [2.3e-21] |
| mafft | -0.21 [3.9e-18] | 0.28 [3.8e-21] | | -0.21 [9.6e-16] | -0.09 [2.6e-07] | -0.20 [1.3e-16] | -0.17 [1.4e-12] |
| dialign | -0.00 [2.2e-13] | 0.50 [2.0e-21] | 0.21 [9.6e-16] | | 0.13 [1.4e-19] | 0.01 [3.4e-14] | 0.04 [2.0e-21] |
| kalign | -0.13 [2.3e-20] | 0.37 [2.4e-21] | 0.09 [2.6e-07] | -0.13 [1.4e-19] | | -0.12 [7.3e-20] | -0.08 [1.0e-11] |
| fsa | -0.01 [5.7e-17] | 0.49 [2.0e-21] | 0.20 [1.3e-16] | -0.01 [3.4e-14] | 0.12 [7.3e-20] | | 0.03 [2.1e-18] |
| clustalo | -0.04 [2.0e-21] | 0.46 [2.3e-21] | 0.17 [1.4e-12] | -0.04 [2.0e-21] | 0.08 [1.0e-11] | -0.03 [2.1e-18] | |

SPS Statistics for paper-data/LINETree-1-L2-R3S-gput3000-mfl2-eval/replicates.csv
- Calculated on all replicate-parameter values

Kruskal-Wallis H-test
H=648.4578156925951, p=8.17573564502539e-137

Wilcoxon signed rank test: mean_diff [p-val]

| | refiner | muscle | mafft | dialign | kalign | fsa | clustalo |
|---|---|---|---|---|---|---|---|
| refiner | | 0.42 [2.0e-21] | 0.08 [3.0e-06] | -0.01 [4.4e-01] | 0.35 [2.0e-21] | -0.00 [5.8e-01] | 0.29 [2.1e-21] |
| muscle | -0.42 [2.0e-21] | | -0.34 [2.0e-21] | -0.43 [2.0e-21] | -0.07 [3.6e-19] | -0.42 [2.0e-21] | -0.13 [2.4e-21] |
| mafft | -0.08 [3.0e-06] | 0.34 [2.0e-21] | | -0.09 [2.3e-08] | 0.27 [2.0e-21] | -0.08 [1.3e-10] | 0.21 [1.5e-19] |
| dialign | 0.01 [4.4e-01] | 0.43 [2.0e-21] | 0.09 [2.3e-08] | | 0.36 [2.0e-21] | 0.01 [6.7e-02] | 0.30 [2.0e-21] |
| kalign | -0.35 [2.0e-21] | 0.07 [3.6e-19] | -0.27 [2.0e-21] | -0.36 [2.0e-21] | | -0.36 [2.0e-21] | -0.06 [7.6e-20] |
| fsa | 0.00 [5.8e-01] | 0.42 [2.0e-21] | 0.08 [1.3e-10] | -0.01 [6.7e-02] | 0.36 [2.0e-21] | | 0.29 [2.0e-21] |
| clustalo | -0.29 [2.1e-21] | 0.13 [2.4e-21] | -0.21 [1.5e-19] | -0.30 [2.0e-21] | 0.06 [7.6e-20] | -0.29 [2.0e-21] | |

SPS Statistics for paper-data/LINETree-1-L2-R3S-eval/replicates.csv
- Calculated on all replicate-parameter values

Kruskal-Wallis H-test
H=157.35787927103888, p=2.1473170450669003e-31

Wilcoxon signed rank test: mean_diff [p-val]

| | refiner | muscle | mafft | dialign | kalign | fsa | clustalo |
|---|---|---|---|---|---|---|---|
| refiner | | 0.19 [2.7e-31] | -0.01 [1.3e-02] | 0.11 [1.8e-30] | 0.21 [9.7e-31] | 0.04 [2.1e-08] | 0.26 [2.7e-31] |
| muscle | -0.19 [2.7e-31] | | -0.21 [2.7e-31] | -0.08 [4.8e-31] | 0.02 [2.7e-02] | -0.16 [2.7e-31] | 0.07 [3.2e-19] |
| mafft | 0.01 [1.3e-02] | 0.21 [2.7e-31] | | 0.13 [5.8e-31] | 0.23 [2.6e-30] | 0.05 [8.7e-04] | 0.28 [2.7e-31] |
| dialign | -0.11 [1.8e-30] | 0.08 [4.8e-31] | -0.13 [5.8e-31] | | 0.10 [1.4e-25] | -0.08 [2.5e-21] | 0.15 [2.7e-31] |
| kalign | -0.21 [9.7e-31] | -0.02 [2.7e-02] | -0.23 [2.6e-30] | -0.10 [1.4e-25] | | -0.18 [1.8e-30] | 0.05 [7.8e-09] |
| fsa | -0.04 [2.1e-08] | 0.16 [2.7e-31] | -0.05 [8.7e-04] | 0.08 [2.5e-21] | 0.18 [1.8e-30] | | 0.23 [2.9e-31] |
| clustalo | -0.26 [2.7e-31] | -0.07 [3.2e-19] | -0.28 [2.7e-31] | -0.15 [2.7e-31] | -0.05 [7.8e-09] | -0.23 [2.9e-31] | |

#### Section S2

SPS Statistics for paper-data/DNATransTree-2-Tigger1-R3S-eval/replicates.csv
- Calculated on all replicate-parameter values
- mean_diff is the difference in mean SPS scores; for example, the mean SPS of refiner is 0.12 higher than the mean SPS of dialign

Kruskal-Wallis H-test
H=120.6690219254254, p=1.1790984491153163e-23

Wilcoxon signed rank test: mean_diff [p-val]

| | refiner | muscle | mafft | dialign | kalign | fsa | clustalo |
|---|---|---|---|---|---|---|---|
| refiner | | 0.19 [1.0e-30] | -0.02 [9.3e-03] | 0.12 [2.7e-29] | 0.19 [4.0e-29] | 0.04 [6.7e-06] | 0.28 [3.0e-30] |
| muscle | -0.19 [1.0e-30] | | -0.21 [4.0e-31] | -0.08 [9.3e-30] | -0.00 [9.3e-02] | -0.16 [2.1e-30] | 0.09 [2.2e-19] |
| mafft | 0.02 [9.3e-03] | 0.21 [4.0e-31] | | 0.14 [1.2e-30] | 0.21 [4.3e-29] | 0.06 [1.3e-02] | 0.30 [3.0e-31] |
| dialign | -0.12 [2.7e-29] | 0.08 [9.3e-30] | -0.14 [1.2e-30] | | 0.07 [3.3e-10] | -0.08 [9.2e-16] | 0.16 [3.2e-30] |
| kalign | -0.19 [4.0e-29] | 0.00 [9.3e-02] | -0.21 [4.3e-29] | -0.07 [3.3e-10] | | -0.15 [3.3e-28] | 0.09 [1.9e-15] |
| fsa | -0.04 [6.7e-06] | 0.16 [2.1e-30] | -0.06 [1.3e-02] | 0.08 [9.2e-16] | 0.15 [3.3e-28] | | 0.24 [4.0e-29] |
| clustalo | -0.28 [3.0e-30] | -0.09 [2.2e-19] | -0.30 [3.0e-31] | -0.16 [3.2e-30] | -0.09 [1.9e-15] | -0.24 [4.0e-29] | |

SPS Statistics for paper-data/DNATransTree-2-Charlie1-R3S-eval/replicates.csv
- Calculated on all replicate-parameter values

Kruskal-Wallis H-test
H=120.10639887175417, p=1.5478548045439774e-23

Wilcoxon signed rank test: mean_diff [p-val]

| | refiner | muscle | mafft | dialign | kalign | fsa | clustalo |
|---|---|---|---|---|---|---|---|
| refiner | | 0.20 [6.8e-31] | -0.03 [1.5e-04] | 0.12 [3.3e-30] | 0.20 [8.7e-30] | 0.04 [7.0e-05] | 0.28 [1.1e-30] |
| muscle | -0.20 [6.8e-31] | | -0.22 [2.7e-31] | -0.07 [2.4e-26] | 0.00 [5.3e-01] | -0.15 [1.8e-30] | 0.08 [7.9e-18] |
| mafft | 0.03 [1.5e-04] | 0.22 [2.7e-31] | | 0.15 [5.8e-31] | 0.23 [3.7e-30] | 0.07 [4.2e-03] | 0.30 [4.5e-31] |
| dialign | -0.12 [3.3e-30] | 0.07 [2.4e-26] | -0.15 [5.8e-31] | | 0.08 [3.4e-13] | -0.08 [7.3e-15] | 0.16 [2.8e-30] |
| kalign | -0.20 [8.7e-30] | -0.00 [5.3e-01] | -0.23 [3.7e-30] | -0.08 [3.4e-13] | | -0.16 [3.0e-28] | 0.08 [2.5e-13] |
| fsa | -0.04 [7.0e-05] | 0.15 [1.8e-30] | -0.07 [4.2e-03] | 0.08 [7.3e-15] | 0.16 [3.0e-28] | | 0.23 [2.8e-28] |
| clustalo | -0.28 [1.1e-30] | -0.08 [7.9e-18] | -0.30 [4.5e-31] | -0.16 [2.8e-30] | -0.08 [2.5e-13] | -0.23 [2.8e-28] | |

SPS Statistics for paper-data/DNATransTree-1-Charlie1-R3S-eval/replicates.csv
- Calculated on all replicate-parameter values

Kruskal-Wallis H-test
H=124.71731977281203, p=1.6621425037307055e-24

Wilcoxon signed rank test: mean_diff [p-val]

| | refiner | muscle | mafft | dialign | kalign | fsa | clustalo |
|---|---|---|---|---|---|---|---|
| refiner | | 0.21 [8.1e-31] | -0.02 [2.9e-03] | 0.13 [6.8e-30] | 0.20 [1.2e-29] | 0.04 [1.5e-05] | 0.28 [2.3e-30] |
| muscle | -0.21 [8.1e-31] | | -0.23 [2.7e-31] | -0.08 [1.4e-28] | -0.01 [1.8e-02] | -0.16 [7.7e-31] | 0.07 [6.7e-15] |
| mafft | 0.02 [2.9e-03] | 0.23 [2.7e-31] | | 0.15 [4.0e-31] | 0.22 [1.8e-29] | 0.07 [1.9e-03] | 0.30 [3.2e-31] |
| dialign | -0.13 [6.8e-30] | 0.08 [1.4e-28] | -0.15 [4.0e-31] | | 0.07 [4.2e-12] | -0.08 [2.1e-15] | 0.16 [1.9e-30] |
| kalign | -0.20 [1.2e-29] | 0.01 [1.8e-02] | -0.22 [1.8e-29] | -0.07 [4.2e-12] | | -0.16 [3.6e-28] | 0.08 [2.7e-13] |
| fsa | -0.04 [1.5e-05] | 0.16 [7.7e-31] | -0.07 [1.9e-03] | 0.08 [2.1e-15] | 0.16 [3.6e-28] | | 0.24 [3.1e-29] |
| clustalo | -0.28 [2.3e-30] | -0.07 [6.7e-15] | -0.30 [3.2e-31] | -0.16 [1.9e-30] | -0.08 [2.7e-13] | -0.24 [3.1e-29] | |

SPS Statistics for paper-data/LINETree-2-L2-R3S-eval/replicates.csv
- Calculated on all replicate-parameter values

Kruskal-Wallis H-test
H=164.8579738273444, p=5.536205077756336e-33

Wilcoxon signed rank test: mean_diff [p-val]

| | refiner | muscle | mafft | dialign | kalign | fsa | clustalo |
|---|---|---|---|---|---|---|---|
| refiner | | 0.14 [2.8e-31] | -0.03 [1.0e-14] | 0.06 [1.5e-29] | 0.15 [2.7e-30] | 0.01 [6.2e-03] | 0.20 [2.7e-31] |
| muscle | -0.14 [2.8e-31] | | -0.17 [4.0e-31] | -0.08 [2.9e-31] | 0.01 [8.9e-02] | -0.13 [2.8e-31] | 0.06 [2.1e-18] |
| mafft | 0.03 [1.0e-14] | 0.17 [4.0e-31] | | 0.09 [1.1e-30] | 0.18 [7.7e-31] | 0.04 [5.2e-06] | 0.23 [2.7e-31] |
| dialign | -0.06 [1.5e-29] | 0.08 [2.9e-31] | -0.09 [1.1e-30] | | 0.09 [1.3e-25] | -0.05 [7.6e-27] | 0.14 [2.7e-31] |
| kalign | -0.15 [2.7e-30] | -0.01 [8.9e-02] | -0.18 [7.7e-31] | -0.09 [1.3e-25] | | -0.14 [1.3e-30] | 0.05 [9.7e-09] |
| fsa | -0.01 [6.2e-03] | 0.13 [2.8e-31] | -0.04 [5.2e-06] | 0.05 [7.6e-27] | 0.14 [1.3e-30] | | 0.19 [2.7e-31] |
| clustalo | -0.20 [2.7e-31] | -0.06 [2.1e-18] | -0.23 [2.7e-31] | -0.14 [2.7e-31] | -0.05 [9.7e-09] | -0.19 [2.7e-31] | |

SPS Statistics for paper-data/LINETree-1-CR1-R3S-eval/replicates.csv
- Calculated on all replicate-parameter values

Kruskal-Wallis H-test
H=152.27827071634064, p=2.5514447811313167e-30

Wilcoxon signed rank test: mean_diff [p-val]

| | refiner | muscle | mafft | dialign | kalign | fsa | clustalo |
|---|---|---|---|---|---|---|---|
| refiner | | 0.18 [2.7e-31] | -0.01 [7.2e-01] | 0.11 [8.9e-31] | 0.18 [1.2e-30] | 0.03 [1.9e-06] | 0.26 [2.7e-31] |
| muscle | -0.18 [2.7e-31] | | -0.20 [2.7e-31] | -0.08 [3.3e-31] | -0.00 [1.4e-01] | -0.16 [2.7e-31] | 0.07 [3.5e-19] |
| mafft | 0.01 [7.2e-01] | 0.20 [2.7e-31] | | 0.12 [2.6e-30] | 0.19 [2.0e-29] | 0.04 [3.4e-02] | 0.27 [2.7e-31] |
| dialign | -0.11 [8.9e-31] | 0.08 [3.3e-31] | -0.12 [2.6e-30] | | 0.07 [2.0e-12] | -0.08 [3.9e-22] | 0.15 [2.7e-31] |
| kalign | -0.18 [1.2e-30] | 0.00 [1.4e-01] | -0.19 [2.0e-29] | -0.07 [2.0e-12] | | -0.15 [3.5e-30] | 0.07 [4.2e-12] |
| fsa | -0.03 [1.9e-06] | 0.16 [2.7e-31] | -0.04 [3.4e-02] | 0.08 [3.9e-22] | 0.15 [3.5e-30] | | 0.23 [3.0e-31] |
| clustalo | -0.26 [2.7e-31] | -0.07 [3.5e-19] | -0.27 [2.7e-31] | -0.15 [2.7e-31] | -0.07 [4.2e-12] | -0.23 [3.0e-31] | |

SPS Statistics for paper-data/LINETree-2-CR1-R3S-eval/replicates.csv
- Calculated on all replicate-parameter values

Kruskal-Wallis H-test
H=159.86521417174743, p=6.324019239237697e-32

Wilcoxon signed rank test: mean_diff [p-val]

| | refiner | muscle | mafft | dialign | kalign | fsa | clustalo |
|---|---|---|---|---|---|---|---|
| refiner | | 0.13 [5.8e-31] | -0.03 [5.4e-10] | 0.06 [6.3e-29] | 0.12 [5.5e-30] | 0.00 [5.0e-01] | 0.19 [2.7e-31] |
| muscle | -0.13 [5.8e-31] | | -0.16 [4.0e-31] | -0.08 [4.2e-31] | -0.01 [7.0e-04] | -0.13 [4.0e-31] | 0.06 [2.7e-16] |
| mafft | 0.03 [5.4e-10] | 0.16 [4.0e-31] | | 0.09 [8.5e-31] | 0.15 [1.6e-29] | 0.03 [3.1e-03] | 0.22 [2.7e-31] |
| dialign | -0.06 [6.3e-29] | 0.08 [4.2e-31] | -0.09 [8.5e-31] | | 0.06 [2.9e-14] | -0.05 [2.1e-26] | 0.14 [2.7e-31] |
| kalign | -0.12 [5.5e-30] | 0.01 [7.0e-04] | -0.15 [1.6e-29] | -0.06 [2.9e-14] | | -0.12 [8.8e-30] | 0.07 [1.2e-13] |
| fsa | -0.00 [5.0e-01] | 0.13 [4.0e-31] | -0.03 [3.1e-03] | 0.05 [2.1e-26] | 0.12 [8.8e-30] | | 0.19 [2.9e-31] |
| clustalo | -0.19 [2.7e-31] | -0.06 [2.7e-16] | -0.22 [2.7e-31] | -0.14 [2.7e-31] | -0.07 [1.2e-13] | -0.19 [2.9e-31] | |

SPS Statistics for paper-data/DNATransTree-1-Tigger1-R3S-gput1500-mfl2-eval/replicates.csv
- Calculated on all replicate-parameter values

Kruskal-Wallis H-test
H=701.8713468214255, p=2.412535022779572e-148

Wilcoxon signed rank test: mean_diff [p-val]

| | refiner | muscle | mafft | dialign | kalign | fsa | clustalo |
|---|---|---|---|---|---|---|---|
| refiner | | 0.69 [2.0e-21] | 0.12 [9.4e-20] | 0.12 [2.0e-21] | 0.37 [2.0e-21] | 0.00 [4.1e-01] | 0.41 [2.0e-21] |
| muscle | -0.69 [2.0e-21] | | -0.57 [2.0e-21] | -0.57 [2.5e-21] | -0.32 [2.2e-21] | -0.69 [2.0e-21] | -0.28 [7.1e-18] |
| mafft | -0.12 [9.4e-20] | 0.57 [2.0e-21] | | -0.00 [6.7e-01] | 0.24 [2.2e-21] | -0.12 [2.0e-21] | 0.28 [4.9e-21] |
| dialign | -0.12 [2.0e-21] | 0.57 [2.5e-21] | 0.00 [6.7e-01] | | 0.25 [3.4e-20] | -0.12 [2.1e-21] | 0.29 [2.0e-21] |
| kalign | -0.37 [2.0e-21] | 0.32 [2.2e-21] | -0.24 [2.2e-21] | -0.25 [3.4e-20] | | -0.37 [2.0e-21] | 0.04 [2.3e-03] |
| fsa | -0.00 [4.1e-01] | 0.69 [2.0e-21] | 0.12 [2.0e-21] | 0.12 [2.1e-21] | 0.37 [2.0e-21] | | 0.40 [2.0e-21] |
| clustalo | -0.41 [2.0e-21] | 0.28 [7.1e-18] | -0.28 [4.9e-21] | -0.29 [2.0e-21] | -0.04 [2.3e-03] | -0.40 [2.0e-21] | |

SPS Statistics for paper-data/LINETree-1-L2-R3S-gput1500-mfl2-eval/replicates.csv
- Calculated on all replicate-parameter values

Kruskal-Wallis H-test
H=688.8965605976895, p=1.5266949844485988e-145

Wilcoxon signed rank test: mean_diff [p-val]

| | refiner | muscle | mafft | dialign | kalign | fsa | clustalo |
|---|---|---|---|---|---|---|---|
| refiner | | 0.71 [2.0e-21] | 0.10 [1.0e-08] | 0.04 [6.3e-08] | 0.36 [2.2e-21] | -0.05 [2.5e-07] | 0.32 [2.0e-21] |
| muscle | -0.71 [2.0e-21] | | -0.62 [2.0e-21] | -0.67 [2.0e-21] | -0.36 [2.0e-21] | -0.76 [2.0e-21] | -0.39 [3.0e-21] |
| mafft | -0.10 [1.0e-08] | 0.62 [2.0e-21] | | -0.06 [6.8e-05] | 0.26 [2.0e-21] | -0.15 [2.0e-21] | 0.22 [6.4e-20] |
| dialign | -0.04 [6.3e-08] | 0.67 [2.0e-21] | 0.06 [6.8e-05] | | 0.32 [2.6e-21] | -0.09 [2.1e-21] | 0.28 [2.0e-21] |
| kalign | -0.36 [2.2e-21] | 0.36 [2.0e-21] | -0.26 [2.0e-21] | -0.32 [2.6e-21] | | -0.41 [2.0e-21] | -0.04 [8.5e-04] |
| fsa | 0.05 [2.5e-07] | 0.76 [2.0e-21] | 0.15 [2.0e-21] | 0.09 [2.1e-21] | 0.41 [2.0e-21] | | 0.37 [2.0e-21] |
| clustalo | -0.32 [2.0e-21] | 0.39 [3.0e-21] | -0.22 [6.4e-20] | -0.28 [2.0e-21] | 0.04 [8.5e-04] | -0.37 [2.0e-21] | |

#### Derived Consensus Sequence Accuracy Metrics

The MSA-derived consensi were evaluated by globally aligning each to the consensus produced by the simulated alignment. Using the alignment of the simulated consensus to itself as the optimal baseline, the fraction of optimal alignment score lost (FOASL) was used to evaluate predicted MSA consensus reconstruction. For each MSA tool, 180 FOASL scores (10 replicates, 18 divergence bins) were collected for analysis. A Kruskal-Wallis H-test was performed to assess if there is a significant difference in the accuracy of reconstructions between MSA tools. Additionally, a Wilcoxon signed rank post-hoc test was performed to generate multiple comparison tables for all pairs of methods.

#### Section S3

Derived Consensus Statistics for paper-data/DNATransTree-1-Tigger1-R3S-eval/replicates.csv
- mean_diff is the difference in mean FOASL scores; for example, the mean FOASL of refiner is 0.25 lower than the mean FOASL of muscle (recall that a lower FOASL score is better)

Kruskal-Wallis H-test
H=106.22090269784896, p=1.2592288564493462e-20

Wilcoxon signed rank test: mean_diff [p-val]

| | muscle | refiner | mafft | dialign | kalign | fsa | clustalo |
|---|---|---|---|---|---|---|---|
| muscle | | 0.25 [2.6e-20] | 0.34 [9.5e-31] | 0.15 [1.8e-20] | 0.01 [9.8e-03] | -0.06 [5.2e-13] | 0.08 [3.0e-13] |
| refiner | -0.25 [2.6e-20] | | 0.09 [1.4e-13] | -0.10 [1.6e-02] | -0.24 [2.1e-14] | -0.31 [6.5e-25] | -0.17 [1.6e-13] |
| mafft | -0.34 [9.5e-31] | -0.09 [1.4e-13] | | -0.19 [8.8e-30] | -0.33 [6.3e-25] | -0.40 [2.7e-31] | -0.26 [1.6e-30] |
| dialign | -0.15 [1.8e-20] | 0.10 [1.6e-02] | 0.19 [8.8e-30] | | -0.14 [2.4e-09] | -0.22 [2.7e-31] | -0.08 [1.3e-11] |
| kalign | -0.01 [9.8e-03] | 0.24 [2.1e-14] | 0.33 [6.3e-25] | 0.14 [2.4e-09] | | -0.07 [1.5e-08] | 0.06 [2.2e-02] |
| fsa | 0.06 [5.2e-13] | 0.31 [6.5e-25] | 0.40 [2.7e-31] | 0.22 [2.7e-31] | 0.07 [1.5e-08] | | 0.14 [1.5e-24] |
| clustalo | -0.08 [3.0e-13] | 0.17 [1.6e-13] | 0.26 [1.6e-30] | 0.08 [1.3e-11] | -0.06 [2.2e-02] | -0.14 [1.5e-24] | |

Derived Consensus Statistics for paper-data/DNATransTree-1-Tigger1-R3S-gput100-mfl2-eval/replicates.csv

Kruskal-Wallis H-test
H=398.4614897652784, p=5.9872753923555094e-83

Wilcoxon signed rank test: mean_diff [p-val]

| | muscle | refiner | mafft | dialign | kalign | fsa | clustalo |
|---|---|---|---|---|---|---|---|
| muscle | | 0.26 [8.9e-20] | 0.02 [5.6e-02] | 0.26 [7.8e-21] | 0.14 [4.4e-08] | -0.35 [3.7e-10] | 0.25 [2.8e-15] |
| refiner | -0.26 [8.9e-20] | | -0.24 [2.5e-14] | 0.00 [4.4e-01] | -0.12 [1.4e-13] | -0.61 [8.0e-21] | -0.01 [3.8e-20] |
| mafft | -0.02 [5.6e-02] | 0.24 [2.5e-14] | | 0.24 [3.7e-14] | 0.12 [2.3e-04] | -0.37 [4.4e-19] | 0.22 [1.3e-08] |
| dialign | -0.26 [7.8e-21] | -0.00 [4.4e-01] | -0.24 [3.7e-14] | | -0.12 [4.5e-14] | -0.61 [3.0e-21] | -0.02 [4.7e-19] |
| kalign | -0.14 [4.4e-08] | 0.12 [1.4e-13] | -0.12 [2.3e-04] | 0.12 [4.5e-14] | | -0.49 [4.7e-20] | 0.10 [8.7e-03] |
| fsa | 0.35 [3.7e-10] | 0.61 [8.0e-21] | 0.37 [4.4e-19] | 0.61 [3.0e-21] | 0.49 [4.7e-20] | | 0.59 [3.0e-16] |

| | | | | | | |
|---|---|---|---|---|---|---|
| clustalo | -0.25 [2.8e-15] | 0.01 [3.8e-20] | -0.22 [1.3e-08] | 0.02 [4.7e-19] | -0.10 [8.7e-03] | -0.59 [3.0e-16] |

Derived Consensus Statistics for paper-data/DNATransTree-1-Tigger1-R3S-gput3000-mfl2-eval/replicates.csv

Kruskal-Wallis H-test

H=442.4205158824569, p=2.0994843534952335e-92

Wilcoxon signed rank test: mean_diff [p-val]

| | muscle | refiner | mafft | dialign | kalign | fsa | clustalo |
|---|---|---|---|---|---|---|---|
| muscle | | 0.68 [2.0e-21] | 0.26 [5.6e-06] | 0.07 [2.0e-01] | -0.12 [4.2e-18] | -0.18 [1.4e-18] | -0.08 [5.0e-09] |
| refiner | -0.68 [2.0e-21] | | -0.42 [1.4e-12] | -0.60 [2.0e-18] | -0.80 [2.0e-21] | -0.86 [2.0e-21] | -0.76 [2.0e-21] |
| mafft | -0.26 [5.6e-06] | 0.42 [1.4e-12] | | -0.18 [3.7e-18] | -0.38 [8.0e-21] | -0.44 [2.0e-21] | -0.34 [4.7e-20] |
| dialign | -0.07 [2.0e-01] | 0.60 [2.0e-18] | 0.18 [3.7e-18] | | -0.20 [7.6e-05] | -0.26 [2.0e-21] | -0.16 [8.5e-04] |
| kalign | 0.12 [4.2e-18] | 0.80 [2.0e-21] | 0.38 [8.0e-21] | 0.20 [7.6e-05] | | -0.06 [9.6e-12] | 0.04 [2.2e-09] |
| fsa | 0.18 [1.4e-18] | 0.86 [2.0e-21] | 0.44 [2.0e-21] | 0.26 [2.0e-21] | 0.06 [9.6e-12] | | 0.10 [1.4e-16] |
| clustalo | 0.08 [5.0e-09] | 0.76 [2.0e-21] | 0.34 [4.7e-20] | 0.16 [8.5e-04] | -0.04 [2.2e-09] | -0.10 [1.4e-16] | |

Derived Consensus Statistics for paper-data/LINETree-1-L2-R3S-gput100-mfl2-eval/replicates.csv

Kruskal-Wallis H-test

H=474.6300611142084, p=2.447215844783801e-99

Wilcoxon signed rank test: mean_diff [p-val]

| | muscle | refiner | mafft | dialign | kalign | fsa | clustalo |
|---|---|---|---|---|---|---|---|
| muscle | | 0.42 [2.0e-21] | 0.11 [1.8e-03] | 0.42 [2.0e-21] | -0.02 [9.2e-01] | -0.27 [2.3e-07] | 0.40 [4.2e-21] |
| refiner | -0.42 [2.0e-21] | | -0.30 [1.4e-15] | 0.00 [8.4e-02] | -0.44 [3.6e-19] | -0.68 [4.3e-21] | -0.01 [1.9e-17] |
| mafft | -0.11 [1.8e-03] | 0.30 [1.4e-15] | | 0.31 [1.6e-15] | -0.13 [1.4e-04] | -0.38 [1.1e-18] | 0.29 [1.4e-12] |
| dialign | -0.42 [2.0e-21] | -0.00 [8.4e-02] | -0.31 [1.6e-15] | | -0.44 [2.3e-19] | -0.68 [3.1e-21] | -0.01 [3.6e-21] |
| kalign | 0.02 [9.2e-01] | 0.44 [3.6e-19] | 0.13 [1.4e-04] | 0.44 [2.3e-19] | | -0.25 [2.4e-09] | 0.43 [3.0e-15] |
| fsa | 0.27 [2.3e-07] | 0.68 [4.3e-21] | 0.38 [1.1e-18] | 0.68 [3.1e-21] | 0.25 [2.4e-09] | | 0.67 [3.3e-18] |
| clustalo | -0.40 [4.2e-21] | 0.01 [1.9e-17] | -0.29 [1.4e-12] | 0.01 [3.6e-21] | -0.43 [3.0e-15] | -0.67 [3.3e-18] | |

Derived Consensus Statistics for paper-data/LINETree-1-L2-R3S-gput3000-mfl2-eval/replicates.csv

Kruskal-Wallis H-test

H=404.25436104899956, p=3.402494769005457e-84

Wilcoxon signed rank test: mean_diff [p-val]

| | muscle | refiner | mafft | dialign | kalign | fsa | clustalo |
|---|---|---|---|---|---|---|---|
| muscle | | 0.42 [2.7e-21] | 0.23 [2.2e-04] | 0.09 [1.1e-01] | -0.14 [2.7e-19] | -0.20 [1.2e-18] | -0.06 [2.1e-05] |
| refiner | -0.42 [2.7e-21] | | -0.19 [5.6e-05] | -0.33 [2.4e-11] | -0.56 [2.0e-21] | -0.62 [2.0e-21] | -0.48 [2.4e-21] |
| mafft | -0.23 [2.2e-04] | 0.19 [5.6e-05] | | -0.13 [2.7e-16] | -0.37 [2.4e-21] | -0.43 [2.0e-21] | -0.29 [1.1e-18] |
| dialign | -0.09 [1.1e-01] | 0.33 [2.4e-11] | 0.13 [2.7e-16] | | -0.23 [3.9e-06] | -0.29 [2.0e-21] | -0.15 [1.4e-03] |
| kalign | 0.14 [2.7e-19] | 0.56 [2.0e-21] | 0.37 [2.4e-21] | 0.23 [3.9e-06] | | -0.06 [3.6e-11] | 0.08 [1.1e-15] |
| fsa | 0.20 [1.2e-18] | 0.62 [2.0e-21] | 0.43 [2.0e-21] | 0.29 [2.0e-21] | 0.06 [3.6e-11] | | 0.14 [2.3e-19] |
| clustalo | 0.06 [2.1e-05] | 0.48 [2.4e-21] | 0.29 [1.1e-18] | 0.15 [1.4e-03] | -0.08 [1.1e-15] | -0.14 [2.3e-19] | |

Derived Consensus Statistics for paper-data/LINETree-1-L2-R3S-eval/replicates.csv

Kruskal-Wallis H-test
H=124.65818545818323, p=1.7104239789294596e-24

Wilcoxon signed rank test: mean_diff [p-val]

| | muscle | refiner | mafft | dialign | kalign | fsa | clustalo |
|---|---|---|---|---|---|---|---|
| muscle | | 0.19 [1.1e-20] | 0.26 [5.7e-31] | 0.10 [1.2e-17] | -0.06 [2.6e-03] | -0.13 [3.7e-28] | 0.04 [8.2e-08] |
| refiner | -0.19 [1.1e-20] | | 0.07 [1.8e-16] | -0.09 [9.8e-01] | -0.25 [1.4e-15] | -0.32 [4.0e-27] | -0.14 [7.3e-15] |
| mafft | -0.26 [5.7e-31] | -0.07 [1.8e-16] | | -0.16 [1.4e-29] | -0.32 [2.4e-29] | -0.39 [5.9e-31] | -0.22 [2.7e-31] |
| dialign | -0.10 [1.2e-17] | 0.09 [9.8e-01] | 0.16 [1.4e-29] | | -0.16 [1.0e-13] | -0.23 [2.8e-31] | -0.05 [1.1e-12] |
| kalign | 0.06 [2.6e-03] | 0.25 [1.4e-15] | 0.32 [2.4e-29] | 0.16 [1.0e-13] | | -0.07 [1.1e-07] | 0.11 [4.5e-07] |
| fsa | 0.13 [3.7e-28] | 0.32 [4.0e-27] | 0.39 [5.9e-31] | 0.23 [2.8e-31] | 0.07 [1.1e-07] | | 0.18 [5.5e-28] |
| clustalo | -0.04 [8.2e-08] | 0.14 [7.3e-15] | 0.22 [2.7e-31] | 0.05 [1.1e-12] | -0.11 [4.5e-07] | -0.18 [5.5e-28] | |

#### Section S4

Derived Consensus Statistics for paper-data/DNATransTree-2-Charlie1-R3S-eval/replicates.csv
- mean_diff is the difference in mean FOASL scores; for example, the mean FOASL of refiner is 0.27 lower than the mean FOASL of muscle (recall that a lower FOASL score is better)

Kruskal-Wallis H-test
H=112.78009265717914, p=5.331986481332645e-22

Wilcoxon signed rank test: mean_diff [p-val]

| | muscle | refiner | mafft | dialign | kalign | fsa | clustalo |
|---|---|---|---|---|---|---|---|
| muscle | | 0.27 [2.4e-20] | 0.35 [1.7e-30] | 0.15 [4.4e-18] | 0.01 [5.8e-02] | -0.08 [6.8e-19] | 0.08 [7.6e-13] |
| refiner | -0.27 [2.4e-20] | | 0.07 [4.2e-11] | -0.13 [8.1e-05] | -0.27 [3.5e-16] | -0.36 [7.1e-27] | -0.20 [7.1e-16] |
| mafft | -0.35 [1.7e-30] | -0.07 [4.2e-11] | | -0.20 [2.1e-30] | -0.34 [2.2e-23] | -0.43 [2.7e-31] | -0.27 [8.9e-31] |
| dialign | -0.15 [4.4e-18] | 0.13 [8.1e-05] | 0.20 [2.1e-30] | | -0.14 [4.0e-08] | -0.23 [2.7e-31] | -0.07 [5.7e-11] |
| kalign | -0.01 [5.8e-02] | 0.27 [3.5e-16] | 0.34 [2.2e-23] | 0.14 [4.0e-08] | | -0.09 [6.7e-10] | 0.07 [3.4e-02] |
| fsa | 0.08 [6.8e-19] | 0.36 [7.1e-27] | 0.43 [2.7e-31] | 0.23 [2.7e-31] | 0.09 [6.7e-10] | | 0.16 [1.1e-27] |
| clustalo | -0.08 [7.6e-13] | 0.20 [7.1e-16] | 0.27 [8.9e-31] | 0.07 [5.7e-11] | -0.07 [3.4e-02] | -0.16 [1.1e-27] | |

Derived Consensus Statistics for paper-data/DNATransTree-1-Charlie1-R3S-eval/replicates.csv

Kruskal-Wallis H-test
H=114.62239274739497, p=2.191115542085094e-22

Wilcoxon signed rank test: mean_diff [p-val]

| | muscle | refiner | mafft | dialign | kalign | fsa | clustalo |
|---|---|---|---|---|---|---|---|
| muscle | | 0.29 [1.1e-22] | 0.36 [1.4e-30] | 0.16 [1.2e-19] | 0.03 [3.1e-04] | -0.07 [3.0e-16] | 0.08 [1.0e-12] |
| refiner | -0.29 [1.1e-22] | | 0.07 [1.2e-11] | -0.13 [3.2e-05] | -0.26 [1.1e-16] | -0.36 [2.0e-27] | -0.21 [6.6e-18] |
| mafft | -0.36 [1.4e-30] | -0.07 [1.2e-11] | | -0.20 [3.1e-30] | -0.33 [1.4e-24] | -0.43 [2.7e-31] | -0.28 [5.2e-31] |
| dialign | -0.16 [1.2e-19] | 0.13 [3.2e-05] | 0.20 [3.1e-30] | | -0.14 [1.9e-08] | -0.23 [2.7e-31] | -0.08 [2.1e-11] |

| | | | | | | | |
|---|---|---|---|---|---|---|---|
| kalign | -0.03 [3.1e-04] | 0.26 [1.1e-16] | 0.33 [1.4e-24] | 0.14 [1.9e-08] | | -0.09 [1.5e-10] | 0.06 [1.1e-01] |
| fsa | 0.07 [3.0e-16] | 0.36 [2.0e-27] | 0.43 [2.7e-31] | 0.23 [2.7e-31] | 0.09 [1.5e-10] | | 0.15 [7.8e-26] |
| clustalo | -0.08 [1.0e-12] | 0.21 [6.6e-18] | 0.28 [5.2e-31] | 0.08 [2.1e-11] | -0.06 [1.1e-01] | -0.15 [7.8e-26] | |

Derived Consensus Statistics for paper-data/DNATransTree-2-Tigger1-R3S-eval/replicates.csv

Kruskal-Wallis H-test

H=102.97441652914945, p=6.006536621566306e-20

Wilcoxon signed rank test: mean_diff [p-val]

| | muscle | refiner | mafft | dialign | kalign | fsa | clustalo |
|---|---|---|---|---|---|---|---|
| muscle | | 0.24 [1.9e-20] | 0.34 [1.9e-30] | 0.15 [6.3e-20] | 0.01 [1.1e-02] | -0.06 [4.9e-11] | 0.08 [4.2e-12] |
| refiner | -0.24 [1.9e-20] | | 0.10 [5.3e-15] | -0.09 [6.1e-03] | -0.23 [5.3e-14] | -0.30 [8.6e-25] | -0.17 [4.7e-13] |
| mafft | -0.34 [1.9e-30] | -0.10 [5.3e-15] | | -0.19 [7.5e-30] | -0.33 [5.1e-25] | -0.40 [2.7e-31] | -0.27 [4.0e-31] |
| dialign | -0.15 [6.3e-20] | 0.09 [6.1e-03] | 0.19 [7.5e-30] | | -0.14 [2.3e-09] | -0.21 [4.0e-31] | -0.08 [5.6e-10] |
| kalign | -0.01 [1.1e-02] | 0.23 [5.3e-14] | 0.33 [5.1e-25] | 0.14 [2.3e-09] | | -0.07 [3.4e-08] | 0.06 [2.9e-02] |
| fsa | 0.06 [4.9e-11] | 0.30 [8.6e-25] | 0.40 [2.7e-31] | 0.21 [4.0e-31] | 0.07 [3.4e-08] | | 0.14 [4.7e-26] |
| clustalo | -0.08 [4.2e-12] | 0.17 [4.7e-13] | 0.27 [4.0e-31] | 0.08 [5.6e-10] | -0.06 [2.9e-02] | -0.14 [4.7e-26] | |

Derived Consensus Statistics for paper-data/DNATransTree-1-Tigger1-R3S-gput1500-mfl2-eval/replicates.csv

Kruskal-Wallis H-test

H=446.9426176834401, p=2.233284271995358e-93

Wilcoxon signed rank test: mean_diff [p-val]

| | muscle | refiner | mafft | dialign | kalign | fsa | clustalo |
|---|---|---|---|---|---|---|---|
| muscle | | 0.81 [2.1e-20] | 0.52 [1.9e-20] | 0.51 [1.8e-20] | -0.11 [1.6e-06] | -0.13 [3.9e-06] | 0.62 [2.4e-21] |
| refiner | -0.81 [2.1e-20] | | -0.29 [3.2e-08] | -0.30 [1.6e-11] | -0.92 [1.0e-19] | -0.94 [1.8e-20] | -0.19 [5.4e-11] |
| mafft | -0.52 [1.9e-20] | 0.29 [3.2e-08] | | -0.01 [8.8e-02] | -0.63 [3.3e-21] | -0.65 [2.0e-21] | 0.10 [3.4e-02] |
| dialign | -0.51 [1.8e-20] | 0.30 [1.6e-11] | 0.01 [8.8e-02] | | -0.62 [3.3e-21] | -0.64 [2.0e-21] | 0.11 [2.5e-03] |
| kalign | 0.11 [1.6e-06] | 0.92 [1.0e-19] | 0.63 [3.3e-21] | 0.62 [3.3e-21] | | -0.02 [1.1e-01] | 0.73 [5.4e-21] |
| fsa | 0.13 [3.9e-06] | 0.94 [1.8e-20] | 0.65 [2.0e-21] | 0.64 [2.0e-21] | 0.02 [1.1e-01] | | 0.75 [2.0e-21] |
| clustalo | -0.62 [2.4e-21] | 0.19 [5.4e-11] | -0.10 [3.4e-02] | -0.11 [2.5e-03] | -0.73 [5.4e-21] | -0.75 [2.0e-21] | |

Derived Consensus Statistics for paper-data/LINETree-1-L2-R3S-gput1500-mfl2-eval/replicates.csv

Kruskal-Wallis H-test

H=449.9610457260423, p=5.004041604318263e-94

Wilcoxon signed rank test: mean_diff [p-val]

| | muscle | refiner | mafft | dialign | kalign | fsa | clustalo |
|---|---|---|---|---|---|---|---|
| muscle | | 0.74 [8.6e-20] | 0.48 [3.0e-19] | 0.61 [3.8e-21] | -0.14 [2.3e-08] | -0.11 [7.4e-05] | 0.68 [2.0e-21] |
| refiner | -0.74 [8.6e-20] | | -0.27 [3.1e-06] | -0.13 [7.2e-03] | -0.88 [4.5e-19] | -0.85 [2.8e-18] | -0.06 [9.6e-02] |
| mafft | -0.48 [3.0e-19] | 0.27 [3.1e-06] | | 0.14 [4.5e-04] | -0.61 [2.0e-21] | -0.59 [2.0e-21] | 0.21 [2.3e-06] |
| dialign | -0.61 [3.8e-21] | 0.13 [7.2e-03] | -0.14 [4.5e-04] | | -0.75 [2.2e-21] | -0.73 [2.0e-21] | 0.07 [2.6e-02] |
| kalign | 0.14 [2.3e-08] | 0.88 [4.5e-19] | 0.61 [2.0e-21] | 0.75 [2.2e-21] | | 0.02 [7.2e-01] | 0.82 [2.1e-21] |
| fsa | 0.11 [7.4e-05] | 0.85 [2.8e-18] | 0.59 [2.0e-21] | 0.73 [2.0e-21] | -0.02 [7.2e-01] | | 0.79 [2.1e-21] |
| clustalo | -0.68 [2.0e-21] | 0.06 [9.6e-02] | -0.21 [2.3e-06] | -0.07 [2.6e-02] | -0.82 [2.1e-21] | -0.79 [2.1e-21] | |

Derived Consensus Statistics for paper-data/LINETree-2-L2-R3S-eval/replicates.csv

Kruskal-Wallis H-test
H=127.75842813965856, p=3.809729567694876e-25

Wilcoxon signed rank test: mean_diff [p-val]

| | muscle | refiner | mafft | dialign | kalign | fsa | clustalo |
|---|---|---|---|---|---|---|---|
| muscle | | 0.05 [1.9e-09] | 0.13 [3.8e-30] | 0.04 [1.4e-11] | -0.10 [4.3e-05] | -0.19 [4.1e-29] | -0.01 [6.8e-01] |
| refiner | -0.05 [1.9e-09] | | 0.09 [2.3e-26] | -0.01 [1.2e-03] | -0.15 [1.0e-09] | -0.24 [8.4e-26] | -0.05 [3.4e-11] |
| mafft | -0.13 [3.8e-30] | -0.09 [2.3e-26] | | -0.10 [4.3e-28] | -0.23 [8.2e-29] | -0.32 [4.9e-31] | -0.14 [2.8e-31] |
| dialign | -0.04 [1.4e-11] | 0.01 [1.2e-03] | 0.10 [4.3e-28] | | -0.13 [2.1e-24] | -0.22 [2.8e-31] | -0.04 [6.5e-17] |
| kalign | 0.10 [4.3e-05] | 0.15 [1.0e-09] | 0.23 [8.2e-29] | 0.13 [2.1e-24] | | -0.09 [1.4e-21] | 0.09 [6.4e-07] |
| fsa | 0.19 [4.1e-29] | 0.24 [8.4e-26] | 0.32 [4.9e-31] | 0.22 [2.8e-31] | 0.09 [1.4e-21] | | 0.18 [9.4e-25] |
| clustalo | 0.01 [6.8e-01] | 0.05 [3.4e-11] | 0.14 [2.8e-31] | 0.04 [6.5e-17] | -0.09 [6.4e-07] | -0.18 [9.4e-25] | |

Derived Consensus Statistics for paper-data/LINETree-1-CR1-R3S-eval/replicates.csv

Kruskal-Wallis H-test
H=113.55915372620962, p=3.6609337522854568e-22

Wilcoxon signed rank test: mean_diff [p-val]

| | muscle | refiner | mafft | dialign | kalign | fsa | clustalo |
|---|---|---|---|---|---|---|---|
| muscle | | 0.18 [2.2e-15] | 0.25 [8.2e-31] | 0.11 [2.8e-22] | -0.03 [9.0e-01] | -0.10 [2.2e-25] | 0.06 [5.5e-11] |
| refiner | -0.18 [2.2e-15] | | 0.08 [1.2e-17] | -0.07 [5.1e-01] | -0.20 [6.7e-08] | -0.28 [8.2e-24] | -0.12 [3.6e-08] |
| mafft | -0.25 [8.2e-31] | -0.08 [1.2e-17] | | -0.14 [2.1e-29] | -0.28 [1.6e-29] | -0.35 [4.2e-31] | -0.20 [2.7e-31] |
| dialign | -0.11 [2.8e-22] | 0.07 [5.1e-01] | 0.14 [2.1e-29] | | -0.13 [2.6e-09] | -0.21 [2.8e-31] | -0.05 [3.2e-15] |
| kalign | 0.03 [9.0e-01] | 0.20 [6.7e-08] | 0.28 [1.6e-29] | 0.13 [2.6e-09] | | -0.08 [8.4e-12] | 0.08 [5.1e-03] |
| fsa | 0.10 [2.2e-25] | 0.28 [8.2e-24] | 0.35 [4.2e-31] | 0.21 [2.8e-31] | 0.08 [8.4e-12] | | 0.16 [1.1e-25] |
| clustalo | -0.06 [5.5e-11] | 0.12 [3.6e-08] | 0.20 [2.7e-31] | 0.05 [3.2e-15] | -0.08 [5.1e-03] | -0.16 [1.1e-25] | |

Derived Consensus Statistics for paper-data/LINETree-2-CR1-R3S-eval/replicates.csv

Kruskal-Wallis H-test
H=130.156952170575, p=1.1911499568528818e-25

Wilcoxon signed rank test: mean_diff [p-val]

| | muscle | refiner | mafft | dialign | kalign | fsa | clustalo |
|---|---|---|---|---|---|---|---|
| muscle | | 0.05 [2.4e-07] | 0.14 [3.0e-30] | 0.06 [1.1e-19] | -0.03 [9.6e-01] | -0.16 [3.1e-29] | 0.02 [1.3e-01] |
| refiner | -0.05 [2.4e-07] | | 0.09 [1.4e-27] | 0.01 [1.1e-06] | -0.09 [8.4e-04] | -0.21 [4.0e-23] | -0.03 [2.8e-07] |
| mafft | -0.14 [3.0e-30] | -0.09 [1.4e-27] | | -0.08 [1.2e-29] | -0.17 [4.7e-29] | -0.29 [4.3e-31] | -0.12 [2.7e-31] |
| dialign | -0.06 [1.1e-19] | -0.01 [1.1e-06] | 0.08 [1.2e-29] | | -0.09 [2.9e-17] | -0.22 [4.1e-31] | -0.04 [7.6e-18] |
| kalign | 0.03 [9.6e-01] | 0.09 [8.4e-04] | 0.17 [4.7e-29] | 0.09 [2.9e-17] | | -0.12 [6.2e-30] | 0.05 [6.4e-03] |
| fsa | 0.16 [3.1e-29] | 0.21 [4.0e-23] | 0.29 [4.3e-31] | 0.22 [4.1e-31] | 0.12 [6.2e-30] | | 0.17 [2.8e-24] |
| clustalo | -0.02 [1.3e-01] | 0.03 [2.8e-07] | 0.12 [2.7e-31] | 0.04 [7.6e-18] | -0.05 [6.4e-03] | -0.17 [2.8e-24] | |

##### Parameters for global alignment of consensi with the Needle alignment tool

Gap Open Penalty: 25
Gap Extension Penalty: 5

Score Matrix:

### FREQS A 0.265 C 0.235 G 0.235 T 0.265

| | A | R | G | C | Y | T | K | M | S | W | N |
|---|---|---|---|---|---|---|---|---|---|---|---|
| A | 9 | 1 | -6 | -15 | -16 | -17 | -12 | -2 | -10 | -4 | -5 |
| R | 1 | 1 | 1 | -15 | -15 | -16 | -6 | -6 | -6 | -7 | -5 |
| G | -6 | 1 | 10 | -15 | -15 | -15 | -2 | -10 | -2 | -10 | -5 |
| C | -15 | -15 | -15 | 10 | 2 | -6 | -9 | -2 | -2 | -9 | -5 |
| Y | -16 | -15 | -15 | 1 | 1 | 1 | -6 | -7 | -7 | -7 | -5 |
| T | -17 | -16 | -15 | -6 | 1 | 9 | -2 | -12 | -11 | -4 | -5 |
| K | -12 | -6 | -2 | -11 | -6 | -2 | -2 | -11 | -7 | -7 | -5 |
| M | -2 | -6 | -10 | -2 | -7 | -12 | -11 | -2 | -7 | -7 | -5 |
| S | -10 | -6 | -2 | -2 | -7 | -11 | -7 | -7 | -2 | -10 | -5 |
| W | -4 | -7 | -10 | -11 | -7 | -4 | -7 | -7 | -10 | -4 | -5 |
| N | -5 | -5 | -5 | -5 | -5 | -5 | -5 | -5 | -5 | -5 | 1 |

**Evaluation with Charlie1 (DNA transposon) and CR1 (LINE)**

Results of the assessment of two additional simulations seeded with the DNA Transposon Charlie1 and the LINE family CR1.

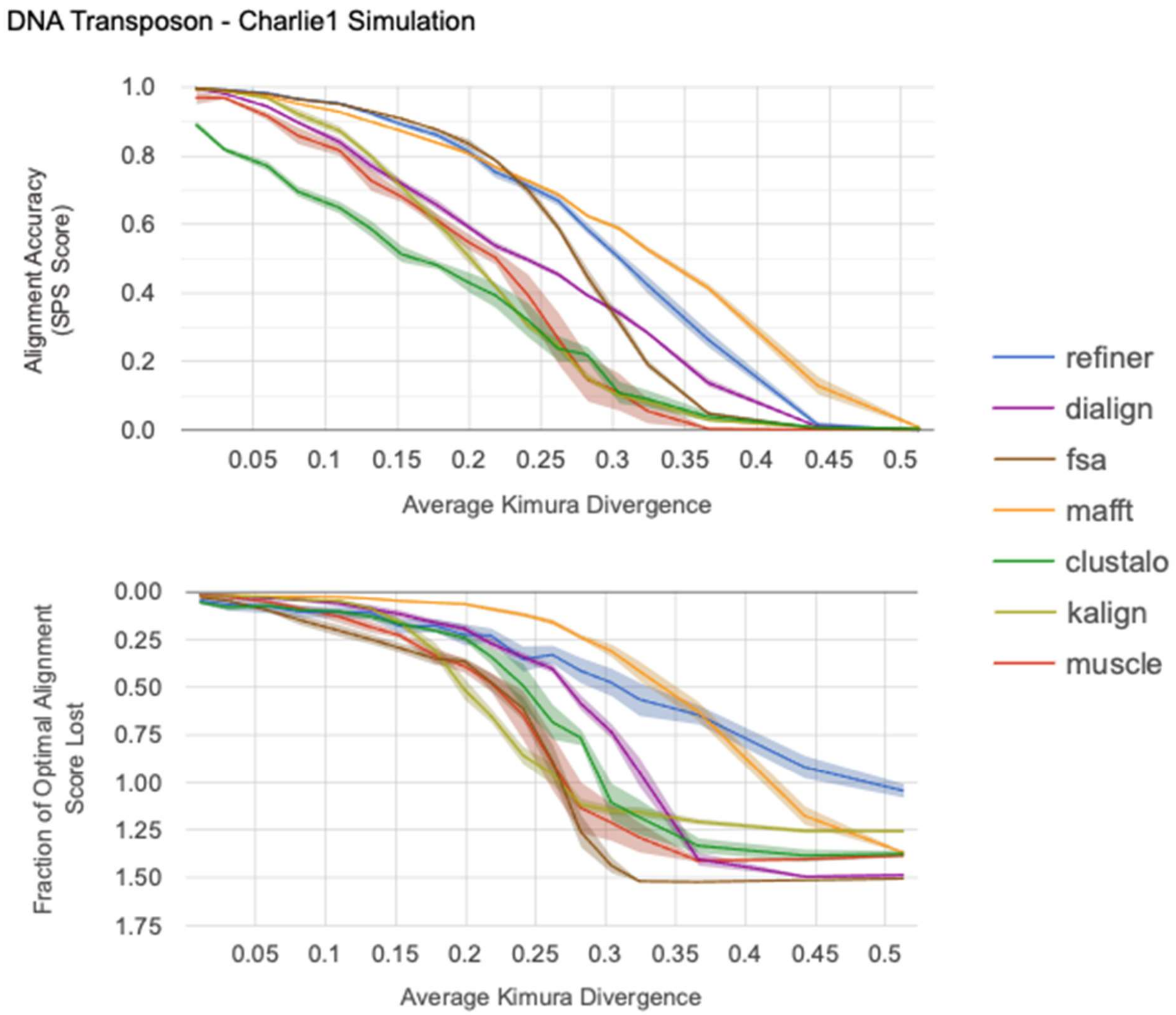

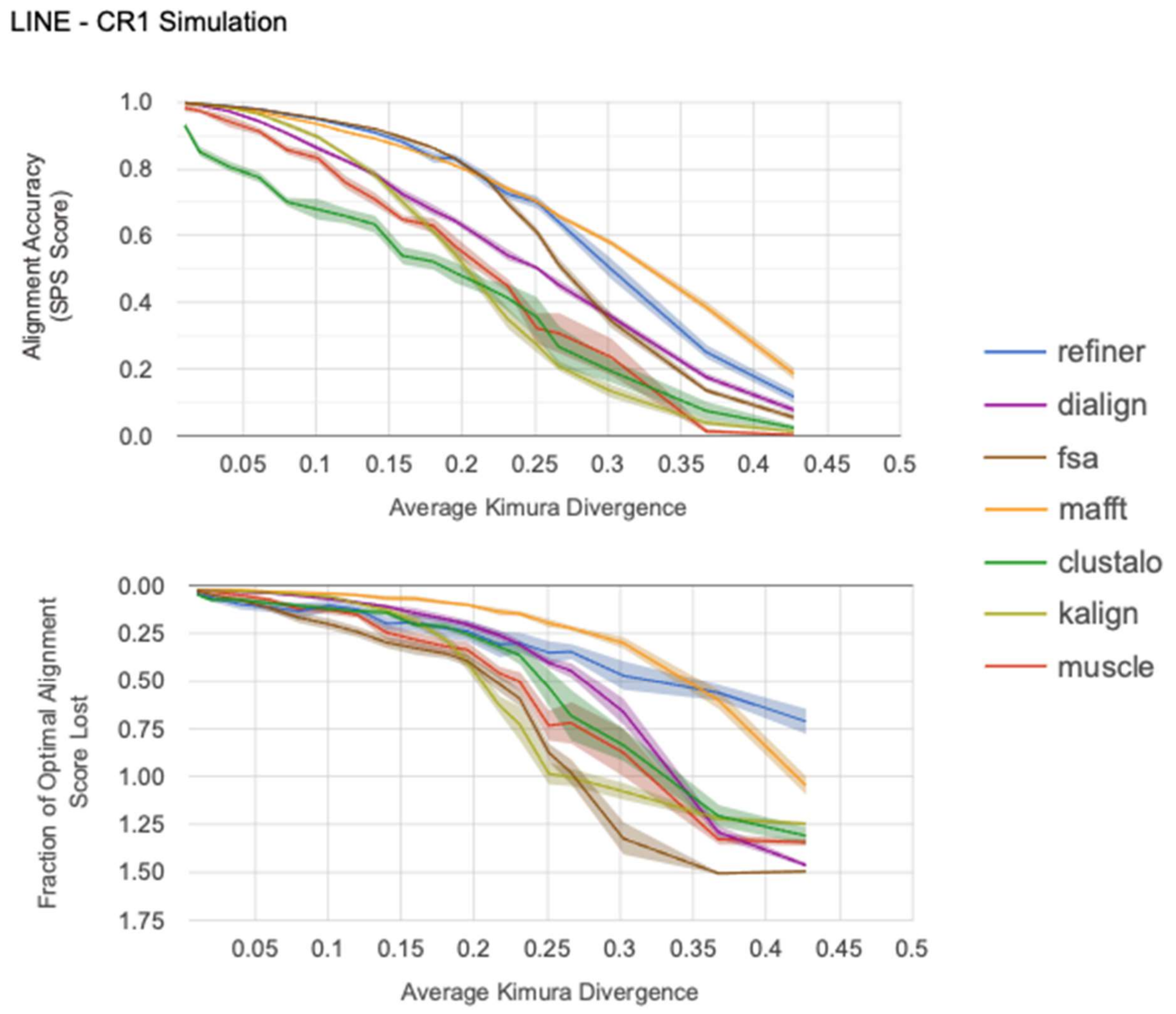

#### Short Regions of Misalignment

Short regions of misalignment are not heavily penalized by the sum of pairs (SPS) scoring metric, but they have the potential to have a profound impact on how the MSA is interpreted. For instance, if short misalignments change the interpretation of sequence occupancy for a given column in the alignment, a derived MSA consensus may be significantly shorter. In supplemental figure S1 the simulated MSA is depicted for a highly fragmented but young DNA transposon family. In figure S2 the predicted alignment from the FSA tool is shown for comparison. In this case the misalignment of a majority of the sequences over a 3 bp stretch causes the consensus caller to consider each of these sequences in the occupancy calculation of the subsequent columns. The large gap between these sequences and the remainder of the residues in the alignment cause a large portion of this MSA to be labeled as insertions.

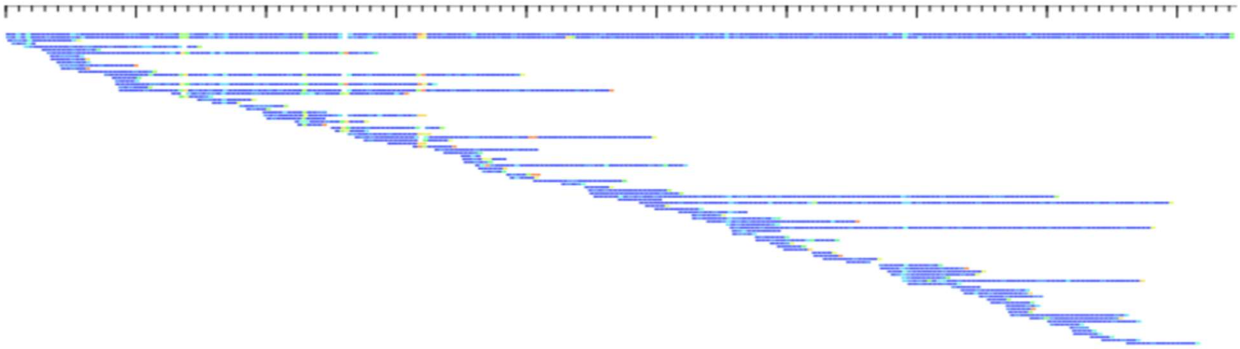

**Figure S1**: Heatmap depiction of a simulated MSA for a young but highly fragmented DNA Transposon family. Cooler colors represent lower divergence of a sequence relative to the MSA consensus over 10bp non-overlapping windows.

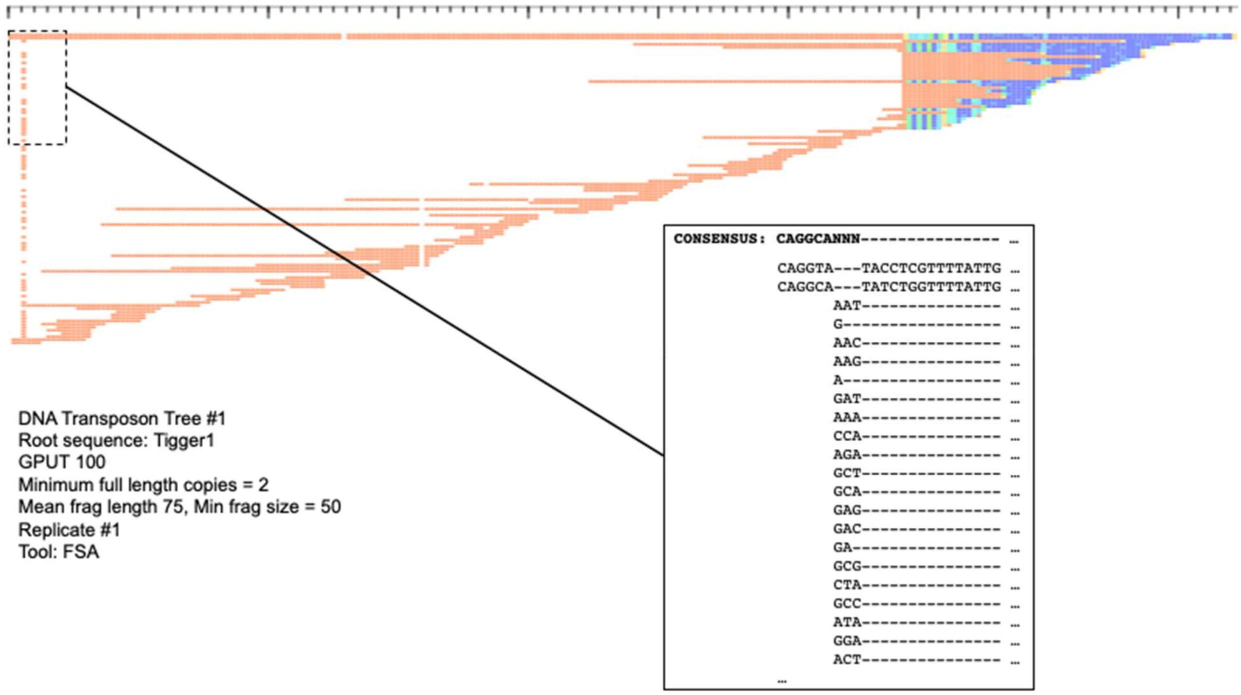

**Figure S2:** Heatmap depiction of the FSA predicted alignment of the simulated sequences from Figure S1. A portion of the alignment from the upper left corner is shown in detail.

#### Mammalian TE Fragmentation Sizes

The fragment size characteristics of 45 mammalian TE families identified in the human HG38 assembly using the RepeatMasker annotation tool.

| Name | Consensus Length | Copies in Genome | Fragments | Longest Fragment (bp) | Mean Fragment Length (bp) | Stdev of Fragment Lengths | Kimura Divergence (ignoring 1 transition at CpG sites) |
|---|---|---|---|---|---|---|---|
| Arthur1 | 3947 | 2135 | 2840 | 2236 | 304.7 | 232.8 | 25.5% |
| Arthur2 | 3700 | 805 | 904 | 1162 | 260.8 | 175 | 33.0% |
| BLACKJACK | 2969 | 1361 | 2037 | 2689 | 414 | 366.9 | 26.2% |
| Charlie1 | 2781 | 1524 | 2025 | 2486 | 279.9 | 296.2 | 21.3% |
| Charlie24 | 2449 | 1211 | 1388 | 2056 | 196.6 | 225.8 | 31.3% |
| Charlie5 | 2624 | 2460 | 3157 | 2604 | 208.7 | 205.1 | 19.6% |
| Charlie7 | 2615 | 2923 | 3946 | 2032 | 275.6 | 218.9 | 28.3% |
| Charlie8 | 2457 | 3214 | 3820 | 2153 | 222.3 | 161.6 | 28.9% |
| Charlie9 | 2787 | 1036 | 1145 | 2373 | 189.5 | 208.1 | 24.2% |
| Cheshire | 2420 | 809 | 1155 | 2412 | 351.4 | 413.7 | 19.1% |
| CR1_Mam | 2204 | 1207 | 1476 | 1233 | 217 | 145.3 | 36.5% |
| EuthAT-2 | 2747 | 855 | 1172 | 2697 | 328.1 | 268.1 | 26.6% |
| FordPrefect | 1683 | 514 | 663 | 1660 | 496.5 | 419.4 | 24.4% |
| hAT-1_Mam | 3469 | 1098 | 1399 | 1797 | 245.5 | 179.5 | 33.9% |
| hAT-5_Mam | 3540 | 606 | 707 | 833 | 205.6 | 132.9 | 32.3% |
| HSMAR2 | 1302 | 1313 | 1870 | 1384 | 609.8 | 450.2 | 12.7% |
| Kanga1 | 779 | 680 | 842 | 1514 | 218.8 | 173.7 | 25.8% |
| L1MA1 | 6302 | 2865 | 4641 | 7805 | 906.5 | 1167.3 | 11.3% |
| L1MA5 | 6300 | 2518 | 4379 | 7202 | 635.7 | 758.2 | 16.2% |
| L1MB1 | 6168 | 3428 | 6482 | 5677 | 512 | 550.9 | 19.2% |
| L1MC1 | 6333 | 7203 | 13849 | 6469 | 573.5 | 585.4 | 17.5% |
| L1MD1 | 6242 | 2662 | 7427 | 5904 | 569.2 | 575.2 | 21.0% |
| L1ME5 | 6194 | 2324 | 3353 | 5559 | 337.3 | 279.9 | 34.5% |
| L1PA10 | 6154 | 5630 | 7997 | 7014 | 962.5 | 1179.3 | 11.5% |
| L1PA5 | 6168 | 9319 | 11892 | 6546 | 1404.3 | 1793.2 | 3.8% |
| L1PB1 | 6151 | 7019 | 13552 | 7673 | 962.7 | 1249.8 | 9.3% |

| Name | Consensus Length | Copies in Genome | Fragments | Longest Fragment (bp) | Mean Fragment Length (bp) | Stdev of Fragment Lengths | Kimura Divergence (ignoring 1 transition at CpG sites) |
|---|---|---|---|---|---|---|---|
| L2-1_AMi | 1235 | 743 | 773 | 771 | 198.5 | 117.3 | 38.2% |
| L2a | 3426 | 76685 | 154946 | 3095 | 270.3 | 239.8 | 35.3% |
| L2b | 3375 | 51557 | 99144 | 2884 | 198.9 | 162.5 | 38.7% |
| L3 | 4099 | 26888 | 41399 | 1752 | 199.7 | 142.4 | 41.1% |
| L4_A_Mam | 4990 | 3924 | 5600 | 1935 | 258.9 | 176.7 | 36.9% |
| L4_C_Mam | 5138 | 2979 | 3976 | 1553 | 222 | 155.3 | 37.5% |
| Looper | 1557 | 356 | 591 | 1559 | 331.3 | 298.6 | 21.2% |
| MamRTE1 | 3812 | 4590 | 5427 | 1447 | 164.7 | 117.4 | 39.2% |
| OldhAT1 | 2842 | 2090 | 2169 | 861 | 95.8 | 76.8 | 34.5% |
| Penelope1_Vert | 1079 | 700 | 725 | 547 | 89.6 | 69.3 | 31.3% |
| Tigger1 | 2418 | 7826 | 13290 | 2490 | 718.8 | 672.3 | 13.0% |
| Tigger10 | 2097 | 1034 | 1234 | 1296 | 221.9 | 159.9 | 35.5% |
| Tigger2 | 2718 | 2516 | 3432 | 2795 | 490.8 | 589.2 | 12.6% |
| Tigger3 | 3029 | 1057 | 1409 | 3027 | 353.4 | 393.7 | 13.0% |
| Tigger7 | 2492 | 2983 | 3354 | 2464 | 221.8 | 198.5 | 13.7% |
| Tigger8 | 667 | 742 | 883 | 650 | 260.2 | 148.9 | 30.0% |
| Zaphod | 4079 | 1618 | 2267 | 3649 | 318.1 | 271.4 | 24.3% |
| Zaphod2 | 3592 | 602 | 778 | 1786 | 223.4 | 221.8 | 24.4% |
| Zaphod3 | 2624 | 909 | 1098 | 1552 | 209.8 | 178.4 | 22.5% |
